# Comparative analysis of macrophage post-translational modifications during intracellular bacterial pathogen infection

**DOI:** 10.1101/2020.05.27.116772

**Authors:** Jeffrey R. Johnson, Trevor Parry, Teresa Repasy, Kristina M. Geiger, Erik Verschueren, Jonathan M. Budzik, David Jimenez-Morales, Billy W. Newton, Emma Powell, Laurent Coscoy, Daniel A. Portnoy, Nevan J. Krogan, Jeffery S. Cox

## Abstract

Macrophages activate robust antimicrobial functions upon engulfing virulent bacteria, yet a wide array of pathogens paradoxically thrive within these innate immune cells. To probe the pathogen-macrophage interface, we used proteomics to comprehensively quantify changes in post-translational modifications (PTMs) of host proteins during infection with three evolutionarily diverse intracellular pathogens: *Mycobacterium tuberculosis, Salmonella enterica* serovar Typhimurium, and *Listeria monocytogenes*. Comparing global phosphorylation and ubiquitylation patterns identified extensive reprogramming of cellular pathways during infection, with ubiquitylation patterns revealing unique pathogen-specific molecular response signatures undetectable by transcriptional profiling. Differential PTM changes during infection with attenuated *M. tuberculosis* cells lacking the ESX-1 virulence determinant revealed extensive modification of phagosome dynamics and antiviral type I interferon activation. We found that *M. tuberculosis*-mediated activation of the antiviral OASL1-IRF7 pathway promotes bacterial replication, uncovering a new mechanism of virus-bacterial synergy. Our data reveals remarkable specificity in innate cellular responses to complex stimuli and provides a resource for deeper understanding of host-pathogen interactions.

## INTRODUCTION

Intracellular bacterial pathogens (IBPs) remain a major cause of morbidity and mortality worldwide, particularly in developing countries (Mitchell et al., 2016). Their ability to grow inside mammalian cells is key for their pathogenesis and present unique challenges for both antibiotic therapies and immune-mediated killing (Kamaruzzaman et al., 2017). A common yet paradoxical feature of many IBPs is their ability to replicate inside macrophages, an innate immune cell type uniquely equipped to directly kill and/or coordinate the elimination of bacteria (Mitchell et al., 2016; Price and Vance, 2014). Indeed, while “life on the inside” provides a niche rich in nutrients, macrophages are protected by overlapping cellular defense mechanisms to sense and destroy intracellular bacteria. Direct killing mechanisms include the inducible generation of reactive oxygen and nitrogen species, antimicrobial peptides, cell-wall degradative enzymes, acidification and fusion of phagosomes with lysosomes, and ubiquitin-targeted autophagy (Price and Vance, 2014). Furthermore, macrophages can undergo dramatic metabolic shifts in response to infection to promote intracellular bacterial killing, although the mechanisms are not entirely clear (Braverman et al., 2016; Galván-Peña and O’Neill, 2014). Likewise, non-cell-autonomous effects of macrophages on inflammatory responses and adaptive immunity include cytokine secretion, cell death, and antigen presentation (Medzhitov, 2008). Understanding how IBPs avert innate and cellular immunity is critical for engineering novel host-directed therapies and vaccination strategies. A comprehensive knowledge of the relationships between intracellular bacterial pathogens and their hosts is fundamental to this approach.

IBPs have historically been grouped into two categories – those that primarily replicate within membrane-bound phagosomal compartments (“vacuolar”, e.g. *Mycobacterium tuberculosis* and *Salmonella enterica* serovar Typhimurium, hereon shortened to *S.* Typhimurium) and those that break free from the initial phagosome and replicate in the macrophage cytosol (e.g. *Listeria monocytogenes*). However, the distinction between cytosolic and vacuolar pathogens has been blurred as it is now appreciated that most vacuolar pathogens, including *M. tuberculosis* and *S.* Typhimurium, also gain access to the cytosol. After phagocytosis, *L. monocytogenes* utilizes its secreted pore-forming toxin LLO to rapidly degrade the phagosomal membrane and replicates in the cytosol (Schnupf and Portnoy, 2007). Both *S.* Typhimurium and *M. tuberculosis* utilize specialized secretion systems (Type III and Type VII, respectively) to permeabilize phagosomal membranes in a much more limited fashion, leading to only a minority of the population patently in the cytosol (Beuzón et al., 2000; Birmingham et al., 2006; Manzanillo et al., 2012; Siméone et al., 2012). Thus, although different pathogens have different strategies for intracellular survival, a common, fundamental requirement appears to be perturbation of intracellular membranes to gain access to the cytosol (Vance et al., 2009).

Cytosolic access, in turn, acts as a pathogen-specific “pattern” that dramatically alters the way macrophages respond to infection compared to non-pathogens (Vance et al., 2009). While a number of pattern recognition receptors (PRRs, e.g. Toll-like receptors, TLRs) are responsible for recognizing microbial products on the macrophage surface or phagosome lumen to primarily activate transcriptional responses, an array of pathways that recognize bacterial products in the cytosol, or membrane damage directly, activate cellular antimicrobial responses including ubiquitin-mediated autophagy, host cell death, and activation of Type I interferon (IFN). Many of these pathways are largely controlled by post-translational modifications (PTMs) of macrophage proteins. For example, autophagy, an ancient anti-microbial pathway that delivers cytosolic pathogens to the lysosome and has pleotropic effects on mammalian immunity (Deretic and Levine, 2018), is regulated by both phosphorylation and ubiquitylation (Herhaus and Dikic, 2015). Pathogens are targeted to autophagy via ubiquitin modification of the damaged phagosome and bacterial cell surface, which recruits autophagy adaptors such as p62 that link ubiquitylated cargo with the autophagy machinery (Huang and Brumell, 2014). Tank Binding Kinase (TBK1) has also been intimately linked with both activation and positive reinforcement of autophagy via phosphorylation of autophagy adaptors during microbial infections (Herhaus and Dikic, 2015). Not surprisingly, IBPs have evolved a variety of mechanisms to subvert or block these modifications to promote their survival (Choy et al., 2012; Mitchell et al., 2018). However, our understanding of all the PTM changes that occur during IBP infection, the cellular pathways they modify, as well as the nature of other types of “patterns” sensed by macrophages, is limited.

Studies of transcriptional responses to individual TLR ligands has demonstrated that only a small amount of specificity can be achieved using different ligands on the same cells (Smale et al., 2014). Likewise, global transcriptional analysis of macrophages infected with different bacterial pathogens also found that while there is some specificity in the responses that can differentiate disparate pathogens with these complex stimuli, the transcriptional response is overwhelmingly monotonic (Nau et al., 2002).

We describe here the results of a quantitative analysis of global ubiquitylation and phosphorylation in macrophages infected with either *M. tuberculosis, S.* Typhimurium, or *L. monocytogenes*. Our analysis uncovered common and divergent pathways modulated by these IBPs that are not governed by transcriptional changes and revealed a high degree of specificity in ubiquitin responses to infection for each pathogen. We have used our results as resource to guide genetic approaches to elucidate a new connection between antiviral and antibacterial responses that controls *M. tuberculosis* growth during infection.

## RESULTS

### Global ubiquitylation analysis reveals common host pathways modulated by intracellular bacterial pathogens

We combined a stable isotope labeling of amino acids in culture (SILAC) approach with K-ε-GG ubiquitin remnant immunoaffinity purification to globally and quantitatively measure changes in host cellular ubiquitylation upon infection with *M. tuberculosis, L. monocytogenes*, and *S.* Typhimurium (Figure 1A) (Kim et al., 2011; Ong et al., 2002; Xu et al., 2010). These pathogens were chosen as they represent three major types of bacteria (acid-fast, gram-positive and gram-negative, respectively) and utilize contrasting pathogenic strategies to replicate inside macrophages. Mouse RAW 264.7 macrophage cells were labeled with light or heavy SILAC medium containing natural L-lysine and L-arginine or ^13^C_6_-L-lysine and ^13^C_6_,^15^N_4_-L-arginine, respectively, and SILAC-labeled cells were either left uninfected or infected with each pathogen in duplicate biological replicates. SILAC labels were inverted between biological replicate experiments to control for SILAC artifacts. Ubiquitylation of host proteins was quantified at two hours post infection for both *L. monocytogenes* and *S.* Typhimurium infections, and six hours post infection for *M. tuberculosis* infection, as these time points represent the peak of ubiquitylation observed around these pathogens in this cell type by immunofluorescence microscopy, consistent with the slower replication of *M. tuberculosis* (Watson et al., 2012). Cells were lysed, proteins were digested with trypsin, and a portion of the resulting peptides were subjected to K-ε-GG ubiquitin remnant immunoaffinity purification. K-ε-GG-enriched peptides were analyzed by high resolution liquid chromatography coupled with liquid chromatography and high-resolution tandem mass spectrometry (LC-MS/MS) analysis. The raw mass spectrometry data were first analyzed by MaxQuant to identify peptides and proteins, localize K-ε-GG modifications, and to extract SILAC ratios (Budzik et al., 2019; Cox and Mann, 2008). The data were subsequently analyzed by the MSstats R package to fit a fixed or mixed effects model of variability at all levels (i.e., variability at the levels of technical replicates, biological replicates, peptides, and proteins), to apply a statistical test to calculate the probability that a given fold-change can be explained by the model, and to adjust probabilities for multiple testing (Choi et al., 2014). The resulting log_2_fold-changes and adjusted p-values for individual ubiquitylation sites in each infection are provided in Table S1.

**Figure 1.**
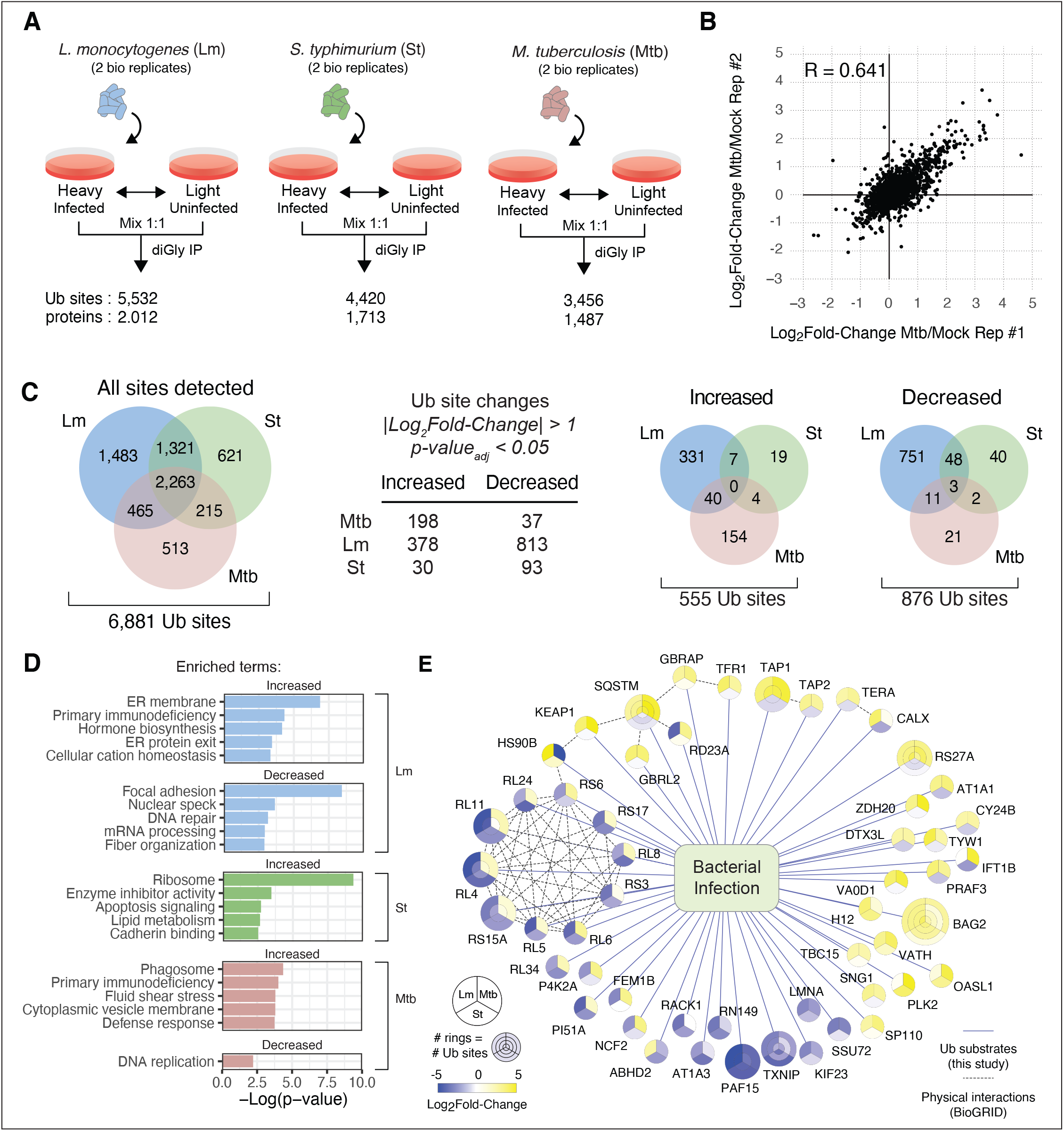
Proteome-wide evaluation of ubiquitylation responses to *L. monocytogenes*, *S. typhimurium*, and *M. tuberculosis* infection. (A) Schematic of experimental design. (B) Representative scatterplot of log_2_fold-changes for ubiquitylation sites quantified in *M. tuberculosis*-infected vs. uninfected cells in two independent biological replicates. (C) Overlap of ubiquitylation sites identified in each infection type (left). Summary of ubiquitylation site changes meeting specified criteria for differential abundance (center). Overlap of differentially increased and decreased ubiquitylation sites by infection type (right). (D) Metascape term enrichment for differentially increased and decreased ubiquitylation sites by infection type. Sites differentially decreased in *S. typhimurium* infection did not yield any significantly enriched terms. (E) Network view of ubiquitylation sites designated as differentially abundant in at least two infection types.

A total of 3,456 ubiquitylation sites on 1,487 proteins were quantified from *M. tuberculosis-*infected cells, 5,532 sites on 2,012 proteins for *L. monocytogenes*-, and 4,420 sites on 1,713 proteins for *S.* Typhimurium-infected cells, representing 6,881 sites in total (Supplementary Table S1). The correlation coefficient between biological replicates for K-ε-GG-containing peptides ranged from 0.428 to 0.516 (Figure 1B; Supplementary Figure S1). 62% of all ubiquitylation sites detected were present in cells infected with at least two bacterial species, and 33% of ubiquitylation sites were quantified in all three (Figure 1C, “All sites detected”). To focus on ubiquitylation events that changed upon infection, we considered sites to be differentially abundant between infection and mock controls if they had a |log_2_fold-change| > 1.0 (i.e., > 2-fold-change) and an adjusted p-value < 0.05. *L. monocytogenes* infection gave rise to the greatest number of ubiquitylation sites that changed significantly, with 378 and 813 sites significantly increased and decreased, respectively (Figure 1C). Interestingly, in contrast to the high overlap in all ubiquitin modified sites detected between macrophages infected with each of the three pathogens, the overlap of the ubiquitylation sites that changed significantly during infection was markedly lower between them (Figure 1C, right Venn diagrams). Using these stringent cutoffs, no ubiquitylation sites were differentially increased by our criteria in all three bacterial infections, while only three ubiquitylation sites were significantly decreased in all three conditions (Figure 1C). Of the three ubiquitylation sites significantly decreased in all three bacterial infections, one site mapped to thioredoxin-interacting protein (TXNIP), a regulator of inflammasomes (Zhou et al., 2011), and two sites mapped to lysine residues 15 and 24 on the PCNA-associated factor of 15 kDa (PAF15). PAF15 is monoubiquitylated at lysine residues 15 and 24 during normal S phase by UHRF1, promoting association with PCNA, and is deubiquitylated under conditions of cell cycle arrest (Karg et al., 2017; Povlsen et al., 2012).

Decreased ubiquitylation of PAF15 at residues 15 and 24 by all three IBPs is consistent with reports that some IBPs induce host cell cycle arrest at the G1/S phase transition (Cumming et al., 2017; Sol et al., 2019).

We performed functional enrichment analysis of significantly regulated ubiquitylation sites using the Metascape tool (Figure 1D) (Tripathi et al., 2015). To compare across the different bacterial infections, the data were filtered for ubiquitylation sites that were detected and quantified in sufficient replicate experiments to calculate log_2_fold-changes and adjusted p-values in all infection conditions. The data were also filtered to remove ubiquitylation sites that did not map unambiguously to a single protein sequence. While some categories are enriched in all three infections, such as trafficking proteins and nucleotide metabolism, and likely represent general host responses to infection, other categories are unique to specific bacterial infections. Ubiquitylation sites that were significantly upregulated by *L. monocytogenes* infection were strongly enriched for integral components of the endoplasmic reticulum (ER) membrane, with the proteins themselves representing a wide range of molecular functions (CANX, DHCR7, TAP1, TAP2, SEC62, BCAP31, EMC8, ARL6IP1, TMCO1, RTN4, HM13; p-value = 1.33×10^−7^). Three of these proteins (DHCR7, BCAP3, RTN4) were also identified as differentially ISGylated proteins during *L. monocytogenes* infection of HeLa cells (Radoshevich et al., 2015). Although di-Gly proteomics cannot distinguish between ubiquitin- and ISG15-modified peptides, our results corroborate those of Radoshevich et al. and strongly suggest that *L. monocytogenes* uniquely interacts with the ER. Interestingly, despite good coverage of ubiquitylated peptides, we observed the fewest number of ubiquitylation changes during *S.* Typhimurium infection, consistent with the notion that this pathogen directly antagonizes the ubiquitin system extensively (Narayanan and Edelmann, 2014). During *M. tuberculosis* infection, sites with increased ubiquitylation were enriched in proteins that shape the phagosome environment, including multiple components of the vacuolar ATPase that functions to acidify the phagosome lumen (ATP6V0D1, ATP6V1H, ATP6V1B2). The extensive modification of v-ATPase is notable as there are only a few reports of its ubiquitylation in the literature and the functional consequences are not well understood (De Luca et al., 2014; Martinez et al., 2017). Likewise, a main subunit (CYBB/gp91phox) of the NADPH oxidase, which creates reactive oxygen species that are both directly antimicrobial and have profound immunoregulatory effects on host cells (Trevelin et al., 2020), is also ubiquitylated during *M. tuberculosis* infection. Likewise, ubiquitylation of mitochondrial and lysosomal proteins is significantly enriched during *M. tuberculosis* infection.

To better visualize the landscape of IBP infection-mediated ubiquitylation changes, we present in Figure 1E a network view of ubiquitylation sites that changed significantly in at least two IBP infections. This visualization illustrates that ubiquitylation sites within the same protein and ubiquitylation sites on different proteins in the sample complex tended to have similar signatures of response to IBP infections. For example, ubiquitylation sites on ribosomal proteins were all observed to decrease in *L. monocytogenes* and *M. tuberculosis* infections but were slightly increased in *S.* Typhimurium infection. Several functional categories stand out, including autophagy, translation, and Type I IFN. For the autophagy receptor p62, also termed sequestosome-1 (SQSTM), all three identified ubiquitylation sites had increased modifications during *L. monocytogenes* and *S.* Typhimurium infections but each decreased during *M. tuberculosis* infection, a similar pattern found in two homologs of ATG8, GABARAP and GABARAP2. As noted above, PAF15 and TXNIP were the only proteins significantly deubiquitylated at the same sites in all three IBP infections and several proteins such as BAG2, a protein recently implicated in autophagy during *M. tuberculosis* infection (Liang et al., 2019), were differentially more ubiquitylated in all three IBP infections though not at exactly the same sites.

Ubiquitin remnant analysis has been previously applied to investigate the host response to *S*. Typhimurium infection in two human cell lines, HCT116 and HeLa (Fiskin et al., 2016). To compare our results with macrophages to this previous study, we converted the mouse ubiquitylation sites to the corresponding human ubiquitylation sites by aligning mouse and human orthologs. If the ubiquitylated residue was aligned to a lysine residue in the human ortholog then the mouse site was converted to the aligned lysine residue in the human protein. Ubiquitylation sites that could not be aligned or did not align to a lysine residue in the corresponding human orthologs were excluded. Pearson correlation coefficients comparing our log_2_fold-change (St-infected/uninfected) profile derived from RAW264.7 cells and the log_2_fold-change (St-infected/uninfected) profiles derived from HCT116 cells at 2 and 5 hours post-infection were 0.192 and 0.185, respectively, while the coefficients comparing log_2_fold-change (St-infected/uninfected) profiles in RAW264.7 cells to HeLa cells at 2 and 5 hours post-infection were 0.262 and 0.248, respectively (Supplementary Figure S2). For comparison, the Pearson correlation coefficient comparing log_2_fold-change (St-infected/uninfected) profiles of HCT116 and HeLa cells at 5 hours post-infection was 0.56. Three proteins were found to be ubiquitylated with a log_2_fold-change > 1 in both RAW264.7 and either HCT116 or HeLa cells: TNF receptor-associated factor 6 (TRAF6, ubiquitin ligase involved in innate immune signaling), CDC42 (actin dynamics), and p62/sequestosome-1 (autophagy). Four proteins were ubiquitylated with a log_2_fold-change < −1 in both RAW264.7 and either HCT116 or HeLa cells: receptor of activated protein C kinase 1 (RACK1), phosphatidylinositol 4-phosphate 5-kinase type-1 alpha, 60S ribosomal protein L34 (RPL34), and PCNA-associated factor (PAF15). Many of the proteins observed with conserved responses in mouse RAW264.7 cells and in human HeLa and/or HCT116 cell lines have been demonstrated to play functional roles in IBP infection and innate immune responses. CDC42 ubiquitylation was demonstrated to lead to linear ubiquitin chain formation and NF-κB activation, TRAF6 ubiquitylation controls the activation of IKK and NF-κB, and p62/Sequestosome-1 ubiquitylation regulates its role in autophagy (Deng et al., 2000; Fiskin et al., 2016; Lamothe et al., 2007; Peng et al., 2017).

### Global phosphorylation analysis identifies pathways commonly and divergently targeted by IBP infection

To identify phosphorylated proteins, we took reserved trypsin-digested material from *L. monocytogenes*-, *S.* Typhimurium-, and *M. tuberculosis*-infected macrophages described above and subjected peptides to phosphopeptide enrichment using Fe^3+^ immobilized metal affinity chromatography (Fe^3+^-IMAC, Figure 2A) (Mertins et al., 2013). Samples were analyzed by LC-MS/MS and analyzed by MaxQuant and MSstats in the same manner as for ubiquitylation analysis described above. Pearson correlations of log_2_fold-change profiles for biological replicates ranged from 0.408 to 0.593 (Supplementary Figure S3). We identified and quantified 9,716 phosphorylation sites in *M. tuberculosis-,* 8,637 sites in *L. monocytogenes-*, and 9,183 sites in *S.* Typhimurium-infected cells, with a total of 11,069 sites identified and quantified in total (Figure 2C; Supplementary Table S2). Phosphorylation sites were considered differentially abundant using the same criteria as for ubiquitylation: |log_2_fold-change| > 1.0 and an adjusted p-value < 0.05. As with ubiquitylation changes, *L. monocytogenes*-infected cells were associated with the greatest number of changes in protein phosphorylation compared to *M. tuberculosis*- and *S.* Typhimurium-infected cells. Of the 11,069 phosphorylation sites that were quantified in total, 74% of sites were quantified in all three IBP infections without missing values. In comparison to ubiquitylation, a higher proportion of phosphorylation sites were differentially abundant during IBP infection. 4% (89 sites) and 15% (239) of phosphorylation sites were differentially increased or decreased, respectively, in all three IBP infections, while zero and 0.3% (3 sites) of ubiquitylation sites were differentially increased or decreased, respectively, in all three IBP infections.

**Figure 2.**
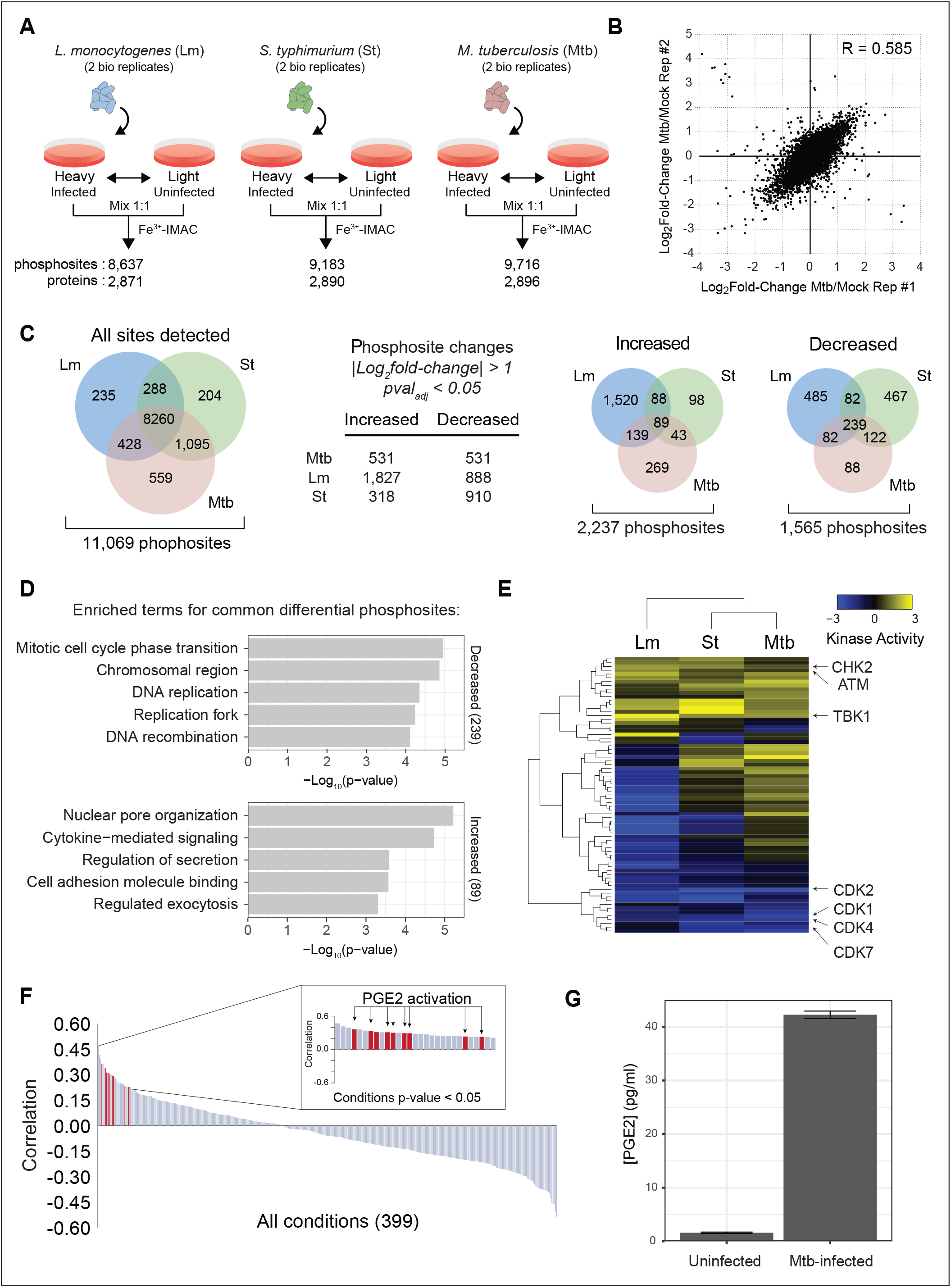
Proteome-wide evaluation of phosphorylation responses to *L. monocytogenes*, *S. typhimurium*, and *M. tuberculosis* infection. (A) Schematic of experimental design. (B) Representative scatterplot of log_2_fold-changes for phosphorylation sites quantified in *M. tuberculosis*-infected vs. uninfected cells in two independent biological replicates. (C) Overlap of phosphorylation sites identified in each infection type (left). Summary of phosphorylation site changes meeting specified criteria for differential abundance (center). Overlap of differentially increased and decreased phosphorylation sites by infection type (right). (D) Metascape term enrichment of phosphorylation sites that were differentially increased and decreased in all infected three types. (E) Heatmap of kinase activities inferred from phosphorylation profiles by PhosFate (Ochoa et al., 2016). Dendrograms indicate the hierarchically clustered Euclidian distances between kinases (left of heatmap) and between infection types (above heatmap). (F) Pearson correlation coefficients comparing the phosphorylation profile of *M. tuberculosis*-infected vs. uninfected cells to profiles within the PhosFate database. Pearson correlations with p-value < 0.05 (inset). (G) PGE2 levels from culture supernatants of *M. tuberculosis*-infected and uninfected cells.

Functional enrichment analysis of the phosphoproteomics data revealed host processes modulated by all three pathogens (Figure 2D). Significant dephosphorylation during infection with each of the three IBPs was found for proteins associated with mitotic cell cycle progression (BRCA1, MCM2, MCM4, NBN, CNOT3, ORC1, CDK14, RB1, RBL1, RRM2, PHF8, ANLN, RCC2, MEPCE; p-value = 1.11×10^−5^) and DNA replication (BRCA1, MCM2, MCM4, NBN, ORC1, RFC1, RRM2, KAT7, WDHD1, SAMHD1, RB1, RBL1, TOP2A, RAD18, DNMT1, H1-3, TSPYL1, RIF1; p-value = 4.37×10^−5^). Proteins of the minichromosome maintenance (MCM) complex are phosphorylated during the transition from G1 to S phase and the downregulation of their phosphorylation by IBP infection is consistent with arrest at this point in the cell cycle (Fei and Xu, 2018). While MCM phosphorylation sites were downregulated by all IBP infections, CDK1 phosphorylation sites at positions T14 and Y15 that are indicative of cell cycle arrest at the G2 to M phase transition were uniquely downregulated by *M. tuberculosis* infection. In contrast, significant increases in phosphorylation in response to all three IBP infections was found for proteins associated with nuclear pore organization (NUP98, TPR, AHCTF1, NUP133, RTN4, NUP35; p-value = 6.06×10^−6^), cytokine-mediated signaling and inflammation (CFL1, EGR1, HNRNPF, IRAK2, JUNB, LCP1, LMNB1, MSN, PRKCD, PSMA5, PSMD1, PSMD2, STX4, TNF, TNFAIP3, TRIM25, SQSTM1, TREX1, TBK1, OTUD4, MAVS, WNK1, TXLNA; p-value = 1.87×10^−5^), and specifically interleukin-1 beta (IL-1β) signaling (EGR1, IRAK2, PSMA5, PSMD1, PSMD2, SQSTM1, TANK, OTUD4; p-value = 1.75×10^−3^). Phosphorylation of nuclear pore components are associated with nuclear envelope breakdown during mitosis, but it is not clear how this relates to bacterial infection (Laurell et al., 2011; Linder et al., 2017). Pro-inflammatory cytokines such as IL-1β produced by monocytes and macrophages are key regulators of the innate immune response to bacterial infection that activates resistance to IBPs, although the mechanisms are not completely understood (van de Veerdonk et al., 2011).

To begin to identify host kinases responsible for the pathogen-dependent phosphorylation changes, we analyzed our data using the PhosFate Profiler tool, which predicts kinase activities from substrate phosphorylation profiles via comparisons with an aggregated database of available known kinase-substrate relationships (Ochoa et al., 2016). This analysis predicted an extensive modulation of host kinases (Figure 2E), seven of which were likely regulated during all three IBP infections including TBK1, which is known to be activated during infection and promotes both IFN production and autophagy targeting (Herhaus and Dikic, 2015). Like TBK1, ATM and CHK2 were also predicted to be upregulated in all three IBP infections, both of which are activated under conditions of DNA damage. Activation of the host DNA damage could be the result of increased oxidative stress generated by NADPH oxidase (Sahan et al., 2018), which itself was ubiquitylated during infection (Figure 1 E, CY24B). Surprisingly, several cyclin-dependent kinases (CDK1, CDK2, CDK4, CDK7) were downregulated with all three IBPs, indicating that infection led to cell cycle arrest. Moreover, kinase activity profiles clustered *S.* Typhimurium and *M. tuberculosis*, which primarily replicate within membrane-delimited phagosomes, more closely together than with *L. monocytogenes*, which patently escapes from the phagolysosomal system and replicates in the cytosol. Kinase activity profile Pearson correlation coefficients were 0.68 for *S.* Typhimurium and *M. tuberculosis*, 0.49 for *S.* Typhimurium and *L. monocytogenes*, and 0.17 for *L. monocytogenes* and *M. tuberculosis*. Although correlative, this analysis will help prioritize targeted experiments to investigate roles for these kinases during infection.

Similar to predicting kinase activity, we used PhosFate to identify published phosphoproteomics profiles that correlated with our data to infer entire pathways modulated by IBP infection. Interestingly, *M. tuberculosis*-infected cells exhibited phosphoproteomics profiles that positively correlated with multiple time points of a phosphoproteomics study of prostaglandin E2 (PGE2) treatment (Figure 2F) (Gerarduzzi et al., 2014). We validated this finding experimentally by measuring PGE2 levels in the culture supernatant of uninfected compared to *M. tuberculosis*-infected cells. PGE2 concentrations were markedly higher in *M. tuberculosis*-infected compared to uninfected cells (Figure 2G). PGE2 is an important regulator of inflammatory responses to *M. tuberculosis* infection (Mayer-Barber et al., 2014) and its expression is stimulated by various cytokines, including IL-1β, and these responses likely drive the correlation between our data of *M. tuberculosis* infection and public data of PGE2 stimulation.

### Ubiquitylation patterns reveal high degree of specificity in innate responses to IBP infection

The ability to distinguish between different types of IBP infection appeared to be a common feature for post-translational modification profiles. Previous work measuring transcriptional outputs have found that an innate immune cell can respond differently when presented with distinct innate immune stimuli, although in most cases these responses were to individual ligands that activate distinct innate receptors rather than complex stimuli such as infection with intact bacteria (Smale et al., 2014). Thus, we sought to compare the level of specificity of PTM changes during infection with that of transcriptional changes. To this end, we also harvested mRNA from macrophages infected with each of the three pathogens and performed RNAseq analysis. We considered mRNA expression to be significantly regulated using the same criteria as for post-translational modifications, |log_2_fold-change|>1.0 and adjusted p-value<0.05. We then compared each type of data (mRNA, phosphorylation, ubiquitylation) among the three different infections using hierarchical clustering based on Euclidian distance (Supplementary Figure S5). While there were some differences in mRNA expression profiles between the three infections, overall the responses were highly similar, with pair-wise Pearson correlation values ranging from 0.78 to 0.87 (Figure 3A). In contrast, phosphorylation profiles between infection were significantly more distinct from one another (R=0.42 - 0.54), and in particular the ubiquitylation profiles were extremely dissimilar (R=0.12 - 0.26, Figure 3A). Both phosphorylation and ubiquitylation analyses clustered *L. monocytogenes* further away from *S.* Typhimurium and *M. tuberculosis* profiles, perhaps a reflection of its different intracellular niche. However, mRNA expression profiles were clustered more closely together than phosphorylation and ubiquitylation profiles, and the *S.* Typhimurium mRNA expression profile clustered more closely with *L. monocytogenes* than with *M. tuberculosis*.

**Figure 3.**
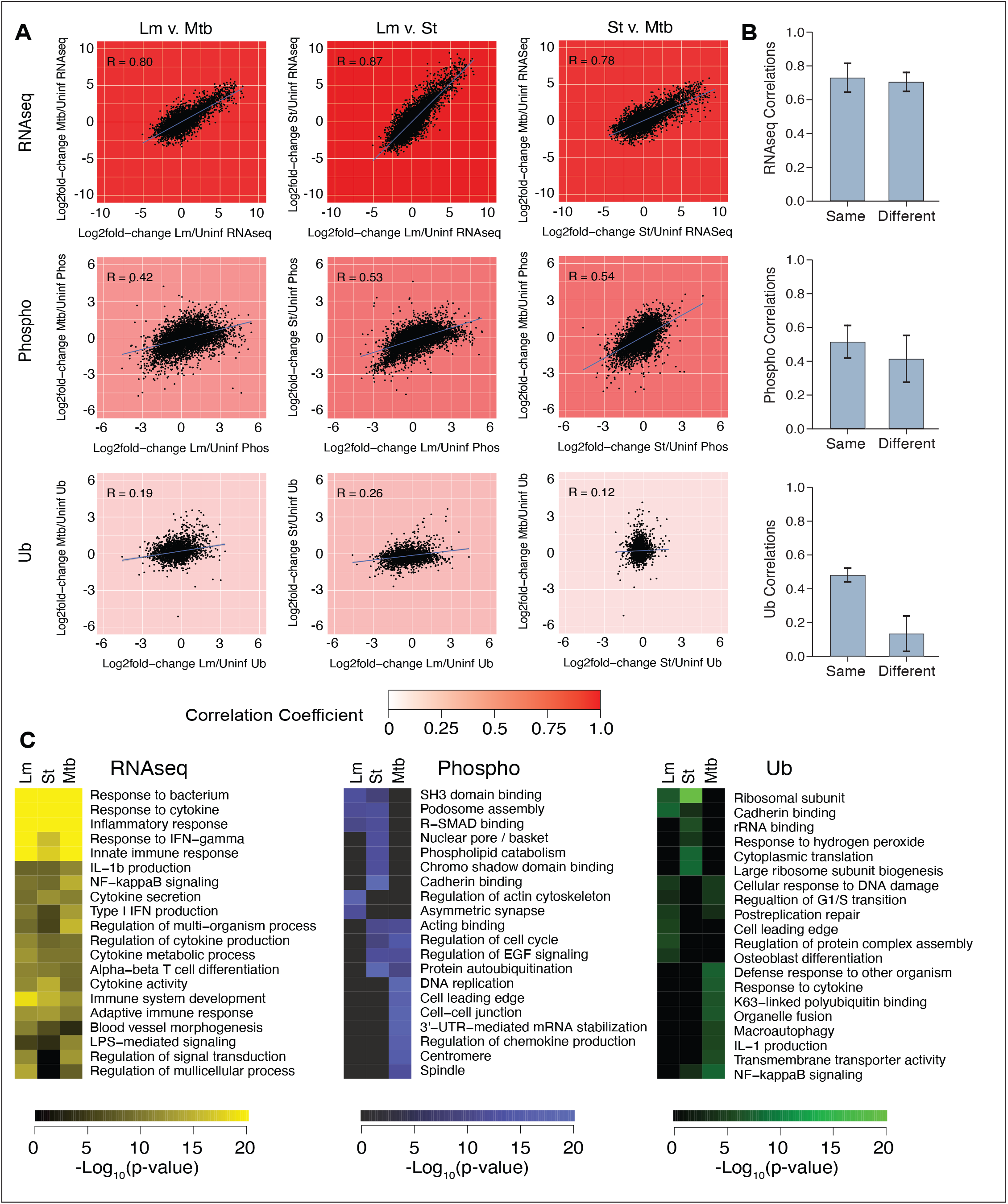
Comparison of genome- and proteome-wide RNA-seq, phosphorylation, and ubiquitylation responses to intracellular bacterial infections. (A) Matrix of scatterplots comparing log_2_fold-changes of indicated infection types for RNA-seq, phosphorylation, and ubiquitylation data types. Matrix cells are shaded white-to-red by Pearson correlation coefficients (upper left corner of each scatterplot). (B) Median and standard deviation of Pearson correlation coefficients of log_2_fold-changes comparing intracellular bacterial infected vs. uninfected cells for biological replicates of the same or different intracellular bacterial infections for RNA-seq, phosphorylation, or ubiquitylation data types. (C) Heatmaps of Metascape terms enriched for differentially abundant RNA-seq, phosphorylation, and ubiquitylation profiles of each intracellular bacterial infection.

Different technologies are associated with different degrees of variability, so we next sought to determine whether the lower correlation coefficients observed for phosphorylation and ubiquitylation profiles were due to biological differences in the data or simply increased technical variability. To test this, we calculated the pairwise Pearson correlation coefficients for each biological replicate of each IBP infection separately for each technology platform (Figure 3B). For mRNA expression and phosphorylation profiles, the correlation coefficients for biological replicate profiles of the same IBP infection were not significantly different from correlation coefficients for different IBP infection profiles, indicating that the decreased correlation of phosphorylation profiles compared to mRNA expression profiles is likely due to increased variability in the technology. However, for the ubiquitylation data, biological replicate profiles were correlated to the same degree as for phosphorylation, but profiles of different IBP infections were significantly lower, indicating that differences between ubiquitylation profiles is not likely due to technical variability. Thus, host ubiquitylation responses indicate that macrophages sense and respond to different infections with a much higher degree of specificity than previously thought.

Pathway-level comparison of functional enrichments also showed that the transcriptional responses are not only more correlated between infections with the different pathogens, but that the enriched pathways themselves are distinct between the different data types (Figure 3C). Indeed, although the raw the phosphoproteomic profiles of log_2_fold-changes were less able to differentiate between IBP infections (Figure 3B), comparison of the public profiles in the PhosFate database revealed a striking ability of condition correlations to identify similar IBP infections (Figure 3C, Supplementary Figure S4). Comparing these correlations between the three IBPs revealed that *S.* Typhimurium and *M. tuberculosis* condition correlation profiles were highly similar (R=0.82), profiles of *L. monocytogenes* and *S.* Typhimurium were less similar (R=0.42), and profiles of *L. monocytogenes* and *M. tuberculosis* were the least similar (R=0.14). In contrast, the phosphoproteomics of *S.* Typhimurium and *M. tuberculosis* log_2_fold-change profiles were the most similar (R=0.57), followed by *S.* Typhimurium and *L. monocytogenes* (R=0.54), and lastly by *L. monocytogenes* and *M. tuberculosis* (R=0.41). Taken together, this analysis also underscores the notion that PTM analysis gives us qualitatively different picture of the cellular response to infection than mRNA abundance.

### *M. tuberculosis* ESX1-dependent ubiquitylation analysis reveals a pattern of host phagosome modulation

The major Type VII secretion system of *M. tuberculosis*, termed ESX-1, is a key virulence pathway that exports bacterial proteins into host macrophages and is essential for the bacterium to replicate inside macrophages (Gröschel et al., 2016). ESX-1 secretion contributes to virulence, at least partially, by permeabilizing host cell phagosome membranes (Watson et al., 2012). To understand the effects of ESX-1 effectors on host cellular pathways, we compared ubiquitylation profiles for cells infected with wild-type and ESX-1 mutant *M. tuberculosis* (Figure 4A). Using the same experimental workflow as above, SILAC-labeled RAW 264.7 macrophage cells were infected with either wild-type or ESX-1 mutant *M. tuberculosis* and compared to uninfected cells in biological duplicates with SILAC labels inverted. At 6 hours post-infection, cells were lysed, total proteins digested with trypsin, and the resulting peptides containing K-ε-GG modifications were enriched and analyzed by LC-MS/MS. Data were analyzed as above with MaxQuant and MSstats. The resulting table of ubiquitylation sites, log_2_fold-changes, and adjusted p-values are provided in Supplementary Table S3. 5,449 ubiquitylation sites on 2,379 proteins were identified in ESX-1 mutant-infected cells, with 67% of sites (3,010) identified in both wild-type and ESX-1 mutant-infected cells (Figure 4B). Approximately 38% of all ubiquitylation sites increased in cells infected with both wild-type and ESX1 mutant *M. tuberculosis* and 27% decreased. To focus on ubiquitylation changes that were ESX1-dependent, we extracted the ubiquitylation sites where |log_2_fold-change|>1 and adjusted p-value < 0.05 in wild-type *M. tuberculosis*-infected cells and where the |log_2_fold-change wild-type:uninfected| - |log_2_fold-change ESX1 mutant:uninfected|>1. These ESX1-dependent ubiquitylation sites are illustrated in network format in Figure 4C. Notably, functional term enrichment analysis of these sites revealed a strong enrichment for proteins associated with the phagosomal membrane, autophagy and IFN signaling (Figure 4D). For example, Perforin-2 (MPEG) is an acid-stimulated cholesterol-dependent cytolysin recently implicated in *M. tuberculosis* resistance that functions on the luminal side of endosomes and activated by ubiquitylation of its C-terminus in the cytosol (McCormack et al., 2015a; 2015b; 2017). We identified this same site in our data, indicating that Perforin-2 is activated in response to cytosolic sensing. Notably, many proteins annotated with lysosome gene ontology were identified with multiple ubiquitylation sites that all increased significantly in wild-type *M. tuberculosis*-infected cells and were significantly less ubiquitylated in ESX1 mutant *M. tuberculosis*-infected cells. The deltex E3 ubiquitin ligase 3L (DTX3L), progranulin (GRN) and vesicle-associated membrane protein 8 (VAMP8) had two ESX1-dependent ubiquitylation sites on each protein, and synaptic vesicle membrane protein VAT-1 homolog (VAT1) had three ESX1-dependent ubiquitylation sites. Also, of note, DTX3L and PARP9 (single ubiquitin modification) mediate mono-ADP-ribosylation of ubiquitin at the C-terminal Gly 76 residue, possibly precluding attachment of ubiquitin to substrate proteins (Yang et al., 2017), and has been implicated in macrophage activation and IFN signaling (Iwata et al., 2016; Zhang et al., 2015). ADP ribosylation of host ubiquitin has been described in *Legionella* infection where the bacterial Sde protein catalyzes the ADP-ribosylation of ubiquitin at residue Arg 42 and subsequent transfer of ubiquitin to a serine residue on the target substrate, Rtn4 (Kotewicz et al., 2017), thus bypassing of the canonical E1-E2-E3 ubiquitin transfer mechanism. It will be of interest in future studies to determine whether DTX3L/PARP9-dependent ADP-ribosylation of ubiquitin catalyzes a similar reaction and plays a role in *M. tuberculosis* pathogenesis.

**Figure 4.**
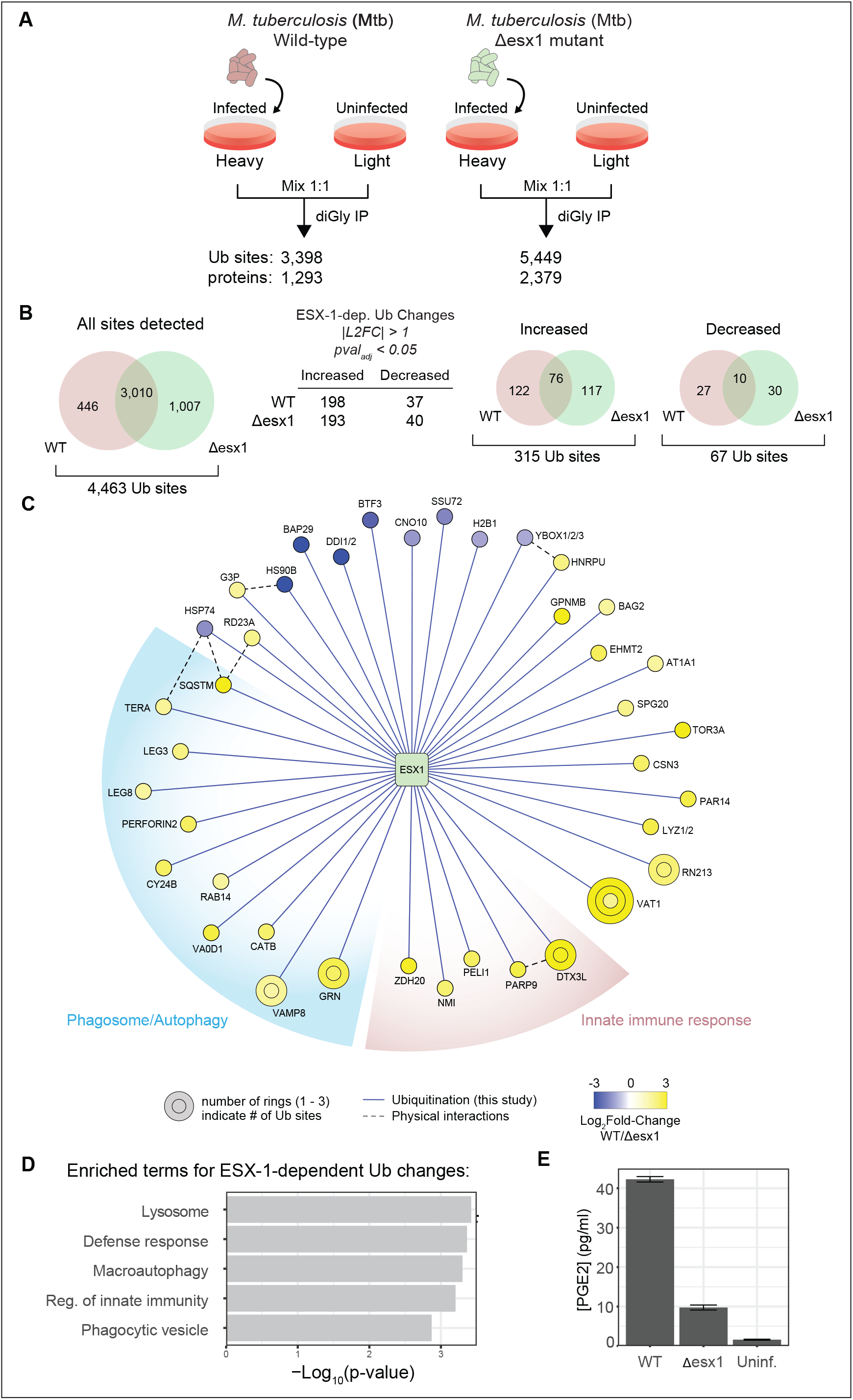
Ubiquitylation analysis of a secretion-defective mutant identifies ubiquitylates components of the lysosome. (A) Schematic of the experimental design. (B) Overlap of ubiquitylation sites detected in wild-type and Δ*esx1 M. tuberculosis*-infected cells (left). Summary of differentially abundant ubiquitylation sites in wild-type and Δ*esx1 M. tuberculosis*-infected cells (center). Overlap of differentially abundant ubiquitylation sites in wild-type and Δesx1 *M. tuberculosis*-infected cells (right). (C) Network view of ESX-1-dependent ubiquitylation changes. (D) Metascape term enrichment for differentially abundant ESX-1-dependent ubiquitylation sites. (E) PGE2 concentrations in cell supernatants for uninfected, WT *M. tuberculosis*-infected, and Δesx1 *M. tuberculosis*-infected cells.

### Overexpression of top ubiquitylated targets identifies IRF7 required for *M. tuberculosis* growth

Based on initial findings from ubiquitylation analysis of *M. tuberculosis*-infected cells, we performed a limited genetic screen to test whether simple overexpression of individual ubiquitylated proteins in macrophages could affect *M. tuberculosis* replication dynamics. 2’,5’-oligoadenylate synthase-like protein 1 (OASL1) was of particular interest because: 1) It was specifically ubiquitylated by wild-type *M. tuberculosis*, 2) we had previously found that *Oasl1* gene transcription was induced in an ESX-1-dependent fashion (Stanley et al., 2007), and 3) OASL1 had not been previously reported to be ubiquitylated. Using an *M. tuberculosis* luciferase reporter strain, we found that over-expression of OASL1 in mouse J774 macrophage cells had the surprising effect of inhibiting *M. tuberculosis* growth over a five-day time course (Figure 5A). Bacterial repression was specific to OASL1, as similar overexpression of either GFP or another highly ubiquitylated protein identified in the di-Gly dataset, Renin Receptor, had no effect on bacterial survival. The observed difference in bacterial growth was not an artifact of premature macrophage death as OASL1-expressing macrophages survived longer than their GFP-expressing counterparts during infection, consistent with decreased bacterial replication. Interestingly, while OASL1 is also ubiquitylated during *S. typhimurium* infection, over-expression of OASL1 had no effect on *S. typhimurium* growth in macrophages (data not shown).

OASL1 has an N-terminal, catalytically-inactive oligoadenylate synthase (OAS) domain, and two ubiquitin-like domains at the C-terminus (Figure 5B). We identified five lysine residues modified by *M. tuberculosis*, one of which, K327, passed our most stringent cutoffs for significance. OASL1 is a translation repressor of approximately 145 genes (Lee et al., 2013), many of which are involved in type-1 interferon, a pathway activated by *M. tuberculosis* during infection (Manzanillo et al., 2012). At steady-state, OASL1 represses translation by binding to 5’ UTR sequence elements in target genes (Lee et al., 2013), but repression is relieved upon viral infection by an unknown mechanism, leading to translation initiation and activation of an antiviral program. We reasoned that ubiquitylation of OASL1 upon infection may lead to destabilization of the protein, but curiously both OASL1 mRNA and protein levels increased in *M. tuberculosis*-infected cells (Figure 5C). Likewise, neither mutation of K327 to arginine nor addition of the proteasome inhibitor MG132 affected OASL1 protein levels, indicating that the activity of OASL1 is inhibited by ubiquitylation (Figure 5D). To determine whether the repressive effect of OASL1 expression on *M. tuberculosis* growth was due to the translational repression of one or more of its targets, we made a three amino acid substitution in OASL1 (R192E, K196E, and K201E), termed OASL1-RKK, that reduces its ability to bind its target mRNAs (Lee et al., 2013). Macrophages expressing OASL1-RKK failed to inhibit bacterial growth (Figure 5E), indicating that the effect of OASL1 on bacterial growth is due to translational repression of one or more of its target transcripts. Of immediate interest was the OASL1 target Interferon Regulatory Factor 7 (IRF7), a transcription factor that is important for sustained type-1 interferon signaling during viral infection (Honda et al., 2005). To test if OASL1 over-expression suppressed bacterial growth exclusively through its effect on IRF7, we co-expressed GFP or IRF7 constructs including its 5’ UTR with or without the OASL1 binding site in J774 cells expressing OASL1. *M. tuberculosis* growth was inhibited in macrophages expressing GFP or an OASL1-sensitive IRF7, but growth was restored in cells expressing an OASL1-insensitive IRF7 (Figure 5F).

**Figure 5.**
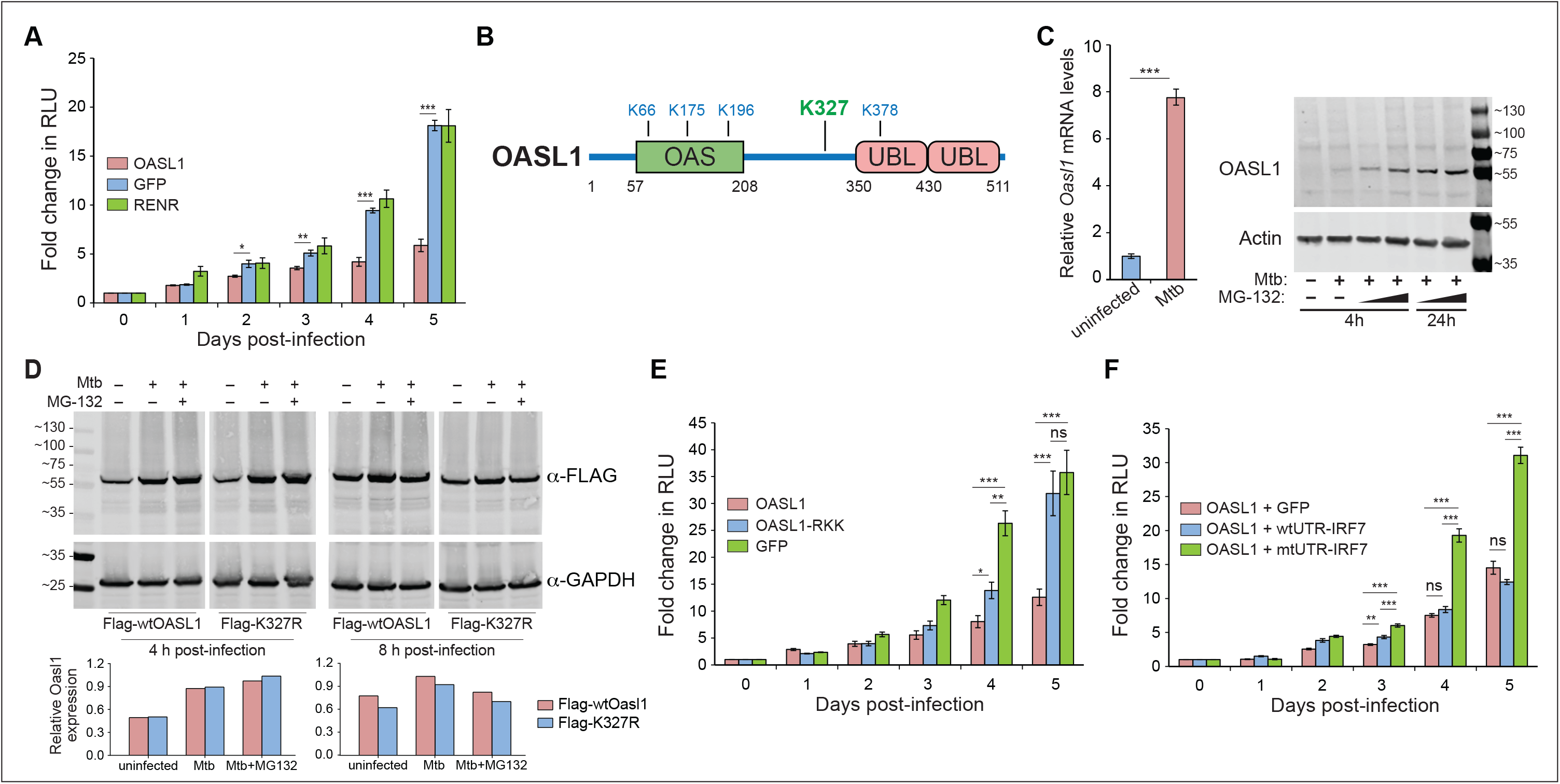
OASL1 restricts *M. tuberculosis* growth by inhibiting translation of IRF7. (A) Bacterial growth assay of J774 mouse macrophages stably overexpressing OASL1, GFP or RenR infected with luminescent *M. tuberculosis* carrying the LuxBCADE reporter operon. Luminescence was measured at indicated time points and values plotted as fold change relative to t=0 (relative luminescent unit, RLU). (B) Diagram of OASL1 showing oligoadenylate synthase (OAS) and Ubiquitin-like (Ubl) domains along with ubiquitylated lysine residues. (C) Oasl1 mRNA levels were assessed by RT-qPCR amplification at six h in mock-infected or *M. tuberculosis*-infected BMMs and data are expressed as a fold increase in expression relative to mock-infected cells. *M. tuberculosis*-infected RAW 264.7 cells treated with MG-132 and levels of OASL1 and β-Actin analyzed by western blot. (numbered lanes 1: uninfected, 2: *M. tuberculosis* only, 3-6: *M. tuberculosis* + MG-132, 3: 50 μM 4 h, 4: 60 μM 4 h, 5: 5 μM 24 h, 6: 10 μM 24 h). (D) *M. tuberculosis*-infected RAW 264.7 cells stably expressing 3XFLAG-Oasl1 (Flag-wtOasl1) or 3XFLAG-Oasl1 with substitution of Lys327 with arginine (Flag-K327R) treated with MG-132 and cell lysates harvested at 4 and 8 h post-infection was analyzed by quantitative western blot and expressed as ratio of FLAG/GAPDH. (E) Bacterial growth assay of J774 cells stably overexpressing GFP, Oasl1 or an RNA-binding deficient Oasl1 mutant (Oasl1-RKK). (F) Bacterial growth assay of J774 cells stably overexpressing Oasl1 and co-selected to overexpress GFP, IRF7 with its wild-type 5’UTR (wtUTR-IRF7) or IRF7 with an eight-base pair mutation in its 5’UTR that disrupts the Oasl1 binding site (mtUTR-IRF7).

This data suggested that IRF7 was a functional target of OASL1 that influences *M. tuberculosis* growth. Indeed, J774 macrophages stably expressing either of two different *Irf7* shRNAs were resistant to bacterial growth relative to controls (Figure 6A), consistent with our observation of the effect of overexpressing OASL1 in these cells. Likewise, CRISPR-mediated homozygous knockout of *Irf7* in RAW 264.7 cells also inhibited *M. tuberculosis* growth (Supplementary Figure S6). Importantly, primary bone marrow-derived macrophages elicited from *Irf7*^−/−^ knockout mice were also highly resistant to *M. tuberculosis* growth (Figure 6B). This effect is unlikely due to activation of known macrophage antibacterial effector mechanisms, as reactive nitrogen species were not induced and trafficking of bacteria to lysosomes, assessed by colocalization with the lysosomal marker ATP6E, remained unaffected in *Irf7*^−/−^ cells (data not shown). When we analyzed the supernatant from infected cells, the bacterial restriction observed from *Irf7*^−/−^ cells was independent of the bacteria’s ability to perforate the phagosome as there were comparable levels of IFN-β and TNF from the supernatant of both wild-type and *Irf7*^−/−^ macrophages (Figure 6C). As expected, while *Irf7^−/−^* macrophages are resistant to *M. tuberculosis* infection, they are highly permissible for the growth of a panel of different viruses (Honda et al., 2005)(Figure 6D and Supplementary Figure S7).

**Figure 6.**
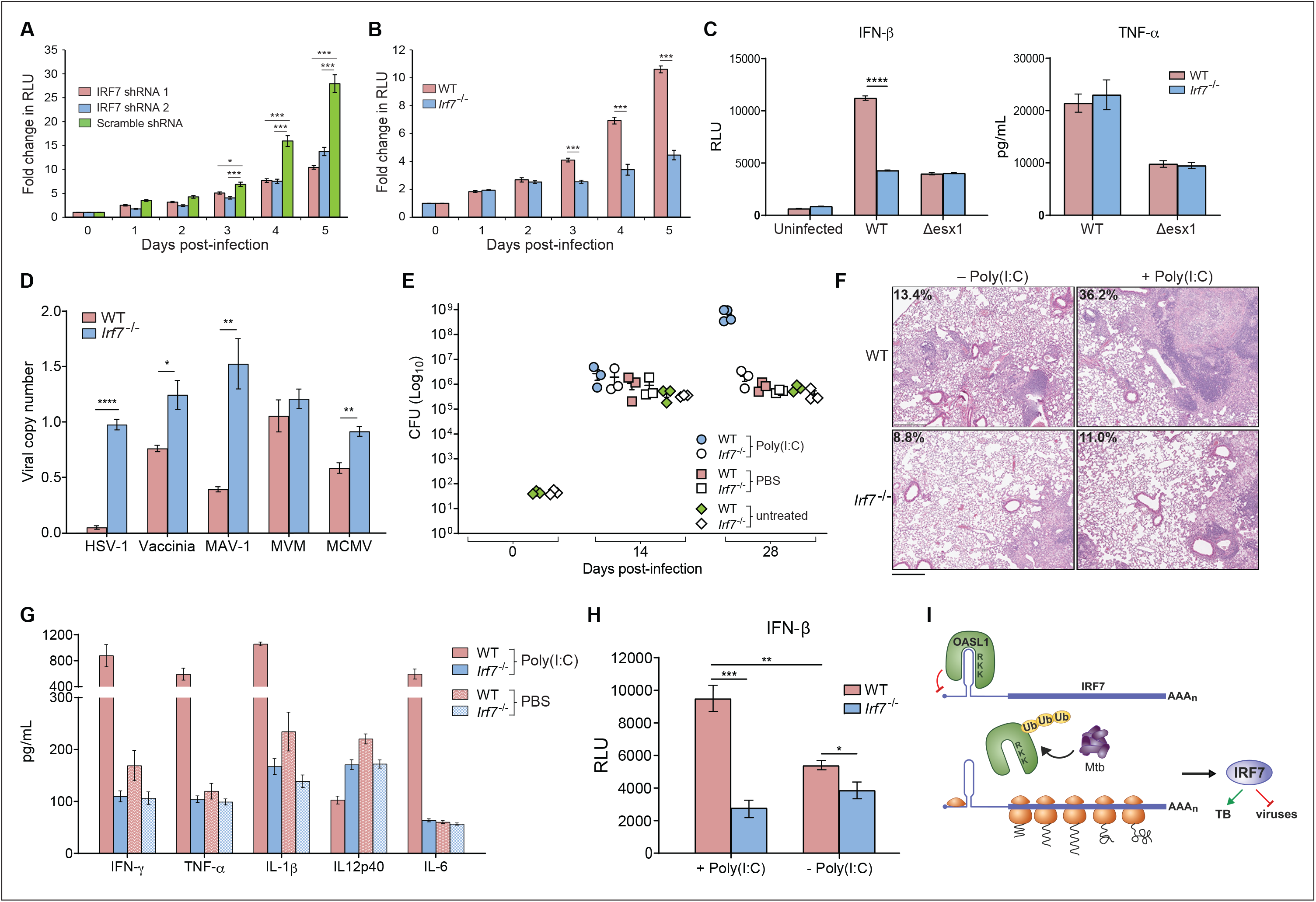
IRF7 promotes *M. tuberculosis* growth ex vivo and in vivo. (A) Bacterial growth assay of stably selected J774 cells transduced with lentiviral constructs expressing shRNAs targeting IRF7 or scrambled shRNA (control). (B) Bacterial growth assay of WT or targeted *Irf7^−/−^* BMMs. (C) BMMs infected with wild-type or Δ*esx1 M. tuberculosis* for 24 h, IFN-β protein levels measured with ISRE-luciferase reporter cells and TNF-α protein levels measured by ELISA. (D) BMMs infected with a panel of viruses for 24 h and viral copy numbers quantified by RT-qPCR in cell culture supernatants. (E) Lung bacterial burdens measured by CFUs of *M. tuberculosis*-infected wild-type and *Irf7^−/−^* mice and treated with poly(I:C) or PBS or untreated at 0, 14 and 28 days post-infection (n=3 or 4 per time point). (F) Representative H&E stained lung tissue sections and percent of pulmonary inflammation of *M. tuberculosis*-infected wild-type and *Irf7^−/−^* mice at 12 weeks post-infection. (n=3, scale baR=0.5 mm) (G) Cytokines measured from cell-free lung lysates of *M. tuberculosis*-infected wild-type or *Irf7^−/−^* mice treated with poly(I:C) or PBS at 28 days post-infection. (H) IFN-β levels measured with reporter cells from lung lysates of *M. tuberculosis*-infected wild-type or *Irf7^−/−^* mice treated with poly(I:C) or PBS at 28 days post-infection. (I) Model of OASL1 in the context of *M. tuberculosis* infection. OASL1 binds to the 5’ UTR region of IRF7 acting as a translational repressor. Upon stimulation with *M. tuberculosis*, OASL1 releases IRF7 transcript permitting the translation of the transcription factor thus initiating downstream Type I interferon response to the pathogen.

To test whether IRF7 plays a role *in vivo* during *M. tuberculosis* infection, we infected wild-type and *Irf7*^−/−^ C57BL/6 mice via the aerosol route and assayed bacterial growth and virulence. Two initial infection experiments performed using mice housed at the UC San Francisco vivarium indicated that *Irf7*^−/−^ mice were resistant to *M. tuberculosis* infection, with a strong inhibition of bacterial growth that resulted in a nearly 10-fold decrease in the stable bacterial burdens in the lung (Supplementary Figures S8A). However, three repeat experiments performed with mice in the UC Berkeley Northwest Animal Facility showed no difference between wild-type and mutant mice, even though the strains of mice and *M. tuberculosis* were identical (Supplementary Figures S8B). Given the role of the gut microbial flora in immune status (Honda and Littman, 2016), we reasoned that microbiota differences between mice housed in the two facilities may have led to increased levels of type I interferon in the mice at UC San Francisco that influenced the phenotype of the *Irf7*^−/−^ mice during *M. tuberculosis* infection. To test this, we sought to provoke tonic type I interferon in mice housed at UC Berkeley by injection of the TLR3 agonist poly(I:C) immediately following infection, which elicits an antiviral response that promotes *M. tuberculosis* infection (Antonelli et al., 2010; Mayer-Barber et al., 2014). As shown in Figure 6E, intraperitoneal injection led to dramatically increased replication in wild-type mice, consistent with previous findings (Antonelli et al., 2010). Importantly, *Irf7*^−/−^ mice were nearly immune to the effects of poly(I:C). The striking difference in bacterial numbers was reflected in the pathological manifestations of TB infection revealed by hematoxylin and eosin staining of lung tissue sections (Figure 6F and Supplementary Figures S9A and S9B) and gross pathology of the spleen (Supplementary Figure S9C). Furthermore, *M. tuberculosis-*infected wild-type mice challenged with poly(I:C) had dramatically higher levels of proinflammatory cytokines IFN-γ, TNF-α, IL-1β, and IL-6 (Figure 6G) along with increased level of IFN-β (Figure 6H) in lung tissues compared to untreated wild-type mice, which only had a slight increase in some of these cytokines (Supplementary Figure S8C). Together, these data demonstrate the antiviral response mediated by IRF7 during *M. tuberculosis* infection promotes exacerbated TB disease in the host.

## DISCUSSION

This work provides a systematic and quantitative analysis of the dynamic changes in ubiquitylation and phosphorylation of the macrophage proteome in response to bacterial infection. Comparisons of these host PTM responses to three different pathogens, each of which parasitize macrophages during normal infection of humans, revealed a dramatic reprogramming of cellular responses during infection, involving both known and unexpected pathways. Likewise, comparisons of PTM changes that are dependent on the key *M. tuberculosis* virulence determinant, ESX-1, have highlighted new proteins and biological pathways providing unparalleled insight into the interactions that underlie pathogenesis of this significant human pathogen.

Importantly, our approach has revealed new cell biological responses that are not captured by transcriptomics. Indeed, over the past 30 years of intense study of the mechanisms of innate immunity, our understanding of macrophage responses to infection has been dominated primarily by studies of transcriptional responses to microbial-derived ligands and PRRs (Medzhitov and Horng, 2009). While gene expression programs certainly play a major role in coordinating a wide variety of inflammatory responses mediated by macrophages, many pathways (e.g. autophagy, metabolism, organelle stress responses) not primarily controlled by changes in transcription are mobilized to fight microbial infection. Our data strongly suggests that PTM analysis is capturing many of these responses.

Likewise, our comparative analysis has identified unique fingerprints of infection for each of the three pathogens, both at the proteome and pathway levels, with ubiquitylation responses in particular showing a high degree of specific responses to each pathogen. This is in sharp contrast to transcriptional responses to infection, which are largely monotonic (Nau et al., 2002; Smale et al., 2014). Although how such specificity in PTM responses is generated remains unknown, we hypothesize that it is reflective of the unique pathogenic strategies of different pathogens. Indeed, the three evolutionarily-distinct bacterial pathogens studied here utilize vastly different virulence factors to mediate interactions with the macrophage, which potentially trigger different cellular responses. For example, the intracellular replicative niche of *L. monocytogenes* is unique in that the bacteria escape from the phagolysosomal system and move by actin-based motility and replicate within the cytosol (Stevens et al., 2006). This may activate a very different repertoire of cellular stress response pathways compared with bacteria that replicate within vacuolar membranes. Our data are consistent with this hypothesis as every comparative analysis we performed identified *L. monocytogenes*-induced PTM profiles as the most distant from those of *S.* Typhimurium and *M. tuberculosis*. Specificity is potentially engendered by integration of overlapping mechanisms of PRR signaling, which recognize bacterial products, and activation of various stress responses, e.g. endoplasmic reticulum stress (Radoshevich et al., 2015), that leads to differential activation of modifying enzymes to reshape the proteome to specifically refine overall cellular responses to match the nature of the invading microbe. It will be interesting to compare PTM responses to other pathogens that also manipulate the actin cytoskeleton, as well as testing mutants of *L. monocytogenes* that are unable to move by actin-based motility.

PTM profiling is a powerful way to generate new hypotheses about host proteins involved in host-pathogen interactions (Fiskin et al., 2016), but the complexity of the many different physiological adaptations of macrophages in response to complex stimuli can make prioritization of targets a daunting task. The use of genetically-defined attenuated bacterial mutants, however, is a simple way to identify changes that are more likely the result of unique pathogenic mechanisms of the pathogen. Indeed, our results comparing the ubiquitylation response of macrophages infected with wild-type or ESX-1^−^ *M. tuberculosis* mutant bacteria are likely the result of physical interactions between the organisms. The ESX-1-specific changes were highly enriched for proteins that are associated with phagosomal membranes, which is permeabilized in an ESX-1-dependent manner and represent a rich source for investigating how macrophages adapt to *M. tuberculosis* infection. For example, the ubiquitylation of vesicle associated membrane protein 8 (VAMP8), a SNARE protein, and the vacuolar ATPase (VA0D1) may represent a new connection between autophagy responses and membrane damage during *M. tuberculosis* infection (Xia et al., 2019). Intriguingly, polymorphisms in the VAMP8 gene has been associated with TB susceptibility in humans (Cheng et al., 2019). Subunits of vATPase have also been recently identified as targets of the Parkin ubiquitin ligase that governs mitophagy (Martinez et al., 2017), providing another potential link between *M. tuberculosis* infection and mitochondrial biology (Härtlova et al., 2018; Manzanillo et al., 2013).

Finally, our finding that ESX-1-dependent activation of the OASL-IRF7 pathway promotes *M. tuberculosis* pathogenesis represents another functional antagonism between antiviral and antibacterial responses. Our previous work on the CBL ubiquitin ligase showed that it also regulates a cell-intrinsic polarization between antiviral and antibacterial immune programs of macrophages but, in contrast to IRF7, promotes antibacterial responses at the cost of antiviral immunity. Antiviral type I interferon responses are detrimental to host resistance to tuberculosis (Ji et al., 2019; Manca et al., 2001), but the mechanisms by which this is mediated, especially the contribution of macrophage-intrinsic effects of IRF7, are not entirely clear. Understanding the mechanism of antiviral and antibacterial antagonism may lead to strategies to break the synergies between *M. tuberculosis* and viral infections such as HIV. Moreover, these results highlight how PTM profiling can facilitate deeper understanding of host-pathogen interactions.

## ACKNOWLEDGEMENTS

This work was funding by NIH grants P01AI063302 (J.S.C and N.J.K.), U19AI106754 (J.S.C and N.J.K.), DP1AI124619 (J.S.C.), U19-AI118610 (J.R.J.)

## AUTHOR CONTRIBUTIONS

Conceptualization: J.S.C. and N.J.K.; Methodology: J.R.J., T.P., T.R., L.C., D.P., N.J.K., and J.S.C.; Software: J.R.J., E.V., and D.J.; Formal Analysis: J.R.J. and E.V.; Investigation: J.R.J., T.P., T.R., K.G., J.B., B.W.N., E.P.; Data Curation: J.R.J.; Writing: J.R.J., T.R., N.J.K., and J.S.C.; Visualization: J.R.J.; Supervision: L.C., D.P., N.J.K., and J.S.C.; Funding Acquisition: D.P., N.J.K., and J.S.C.

## METHODS

### Ethics Statement

Animal experiments were conducted in strict accordance with animal use protocol (AUP-2015-11-8096) approved by the Animal Care and Use Committee at University of California, Berkeley, in adherence with federal regulations provided by the National Research Council and National Institutes of Health.

### Mice and macrophages

*Irf7*^−/−^ mice on the C57BL/6 background were provided by from Gregory Barton at University of California Berkeley. Wild-type C57BL/6J mice were purchased from Jackson Laboratories. Unless otherwise noted, all mice were bred in-housed at either University of California Berkeley or University of California San Francisco. All mice experiments adhered strictly to the regulatory standards set forth by Institutional Animal Care and Use Committee at University of California Berkeley. Bone marrow was isolated from mouse femurs of 8-12-week-old mice. Cells were cultured for seven days in DMEM supplemented with 20% fetal bovine serum, 2 mM L-glutamine and 10% conditioned media derived from 3T3-MCSF cells. J774A.1 (ATCC TIB-67) and RAW 264.7 (ATCC TIB-71) mouse macrophage cell lines were obtained from ATCC.

### Bacterial strains

The following bacterial strains were used: wild-type and Δ*eccD M. tuberculosis* (Erdman), wild-type *L. monocytogenes* (10403s) and wild-type *S. typhimurium* (SL1344). Mycobacterial strains were thawed from frozen stock aliquot and cultured in Middlebrook 7H9 broth (BD #271310) with 10% Middlebrook OADC Enrichment (BD #212351), 0.5% glycerol and 0.05% Tween 80 in roller bottles at 37°C until log phase. For bacterial growth assays measured by luminescence, *M. tuberculosis* expressing codon-optimized *luxBCADE* was used.

### Plasmids and Reagents

pENTR1A no ccdB (w48-1), pLenti CMV Puro DEST (w118-1) and pLenti CMV Blast DEST (706-1) were gifts from Eric Campeau & Paul Kaufman (Addgene plasmids #17398, #17452, #17451). pMD2.G and psPAX2 were gifts from Didier Trono (Addgene plasmids #12259, #12260). shRNA constructs were cloned into the Mission pLKO.1 lentivirus system from Sigma. The following antibodies were used: anti-Oasl1 (Abcam ab116220), anti-ubiquitin (Millipore #04-263), anti-FLAG M2 (Sigma #F1804), anti-GAPDH (Sigma #G9545), anti-β-Actin (Santa Cruz #47778). The following reagents were used: MG-132 (Sigma #474790) and poly(I:C) (HMW) (InvivoGen #tlrl-pic).

### Lentiviral overexpression and knockdown

ORFs were cloned into the pENTR1A construct and transferred into the pLENTI CMV puro or blast vectors using a Gateway enzyme mix (Invitrogen #56484). Lentivirus was produced by co-transfection of the pLenti CMV plasmids with pMD2.G and psPAX2 plasmids. Lentivirus expressing shRNAs targeting mouse *Irf7* transcripts (*Irf7* shRNA1, Sigma TRCN0000077292 and *Irf7* shRNA2, Sigma TRCN0000077289) were generated using the Mission PLKO.1 lentivirus system from Sigma. A lentivirus expressing a scrambled, non-targeting shRNA was used as a control. J774 cells were transduced and selected on Puromycin (2 μg/mL) or Blasticidin (0.75 μg/mL).

### Macrophage infection with *M. tuberculosis*

For macrophage infections measuring intracellular bacterial growth, cultures of *M. tuberculosis* expressing a codon-optimized *luxBCADE* operon was grown to logarithmic phase and washed twice with PBS, sonicated to disperse clumps, and diluted into DMEM supplemented with 10% horse serum. Media was removed from macrophages; monolayers were overlaid with the bacterial suspension at MOI of one or two and centrifuged for 10 minutes at 162*g*. After centrifugation, the monolayers were washed twice with PBS and bacterial luminescence was read on a Tecan Infinite 200 microplate reader or SpectraMax L Microplate Reader (Molecular Devices). At indicated time points, bacterial luminescence was determined as described previously (Penn et al., 2018).

### Viral infection of Macrophages

Primary WT and IRF7 KO BMMs were plated at 200,000 cells per well in 24-well plate. Cells were inoculated with a panel of viruses at MOI of 10 and spun at 750xg for 2 hours. Supernatant was removed and cells were washed 3 times before adding fresh media. After 24 hours, infected cell supernatant was collected and viral DNA was extracted (DNeasy Blood & Tissue Kit, Qiagen 69504). qPCR was performed on processed viral DNA samples to determine viral copy number.

### Western blots

Protein concentration from cell lysates were quantified by BCA (Pierce BCA Protein Assay Kit, Thermo Scientific). Protein lysates were obtained by lysing cells in 0.1% SDS and protease inhibitor cocktail tablets (Roche #04693124001). Cell lysates were separated by SDS-PAGE (Bio-Rad 4-20% Mini-PROTEAN TGX Precast Gel) and proteins were transferred to nitrocellulose membranes. Odyssey blocking buffer was used to block the membranes and indicated antibodies were used for probing. After incubation with IRDye secondary antibodies, Odyssey scanner (LI-COR) was used to image the membrane.

### RNA-Seq

For RNA-seq, three independent experiments were performed. For mock infections, cells were treated identical to infected cells less the addition of the pathogen. Bone marrow-derived macrophages were seeded at a density of 2.5e6 cells per well in four-well cell culture treated plates 24 hours prior and infected either *M. tuberculosis*, *L. monocytogenes* or *S. typhimurium* at MOI 10. At 2 and 6 h post infection, infected and uninfected cells (mock infected) were washed with PBS and lysed with TRIzol Reagent (Invitrogen). After phase separation produced by the addition of chloroform, equal volume of 70% ethanol was added to the aqueous phase. RNA was isolated using Invitrogen PureLink RNA Mini Kit following manufacturer’s instructions and treated with DNase (New England Biolabs). Biological triplicate samples were submitted to DNA Technologies and Expression Analysis Core at UC Davis Genome Center for the generation of 3’Tag RNA-Seq libraries. The Bioinformatics Core at UC Davis performed the differential gene expression analysis.

### Mouse infections and histology

Co-housed wild-type and *Irf7^−/−^* C57BL/6 mice (8-12 week old, male and female) were inoculated with low dose of *M. tuberculosis* via aerosol route using the Inhalation Exposure System by Glas-Col (UCB mouse infections) or Madison chamber device (UCSF mouse infections). One day after infection, infected mice were euthanized and the entire lung was homogenized and plated on 7H10 plates supplemented with 10% OADC and 0.5% glycerol to determine the delivered dosage. CFU were enumerated approximately 21 days after plating. Infected mice were separated into three groups; group one received 5.0 mg/kg of poly(I:C) via intraperitoneal injection, group two received comparable volume of PBS via IP (based on body weight as group one) and group three received no treatment. Mice from group one and two were weighed prior to each injection and mice were injected three times a week starting at one day post-infection. At indicated time points, wild-type and *Irf7^−/−^* mice (age and sex matched) were euthanized and lungs lobes were excised and either homogenized for CFU by plating on 7H10 plates or fixed with 10% buffered formalin. Fixed tissues were paraffin embedded and 5 μm sections were stained with hematoxylin and eosin.

### Cytokine measurements

Measurement of IFN-β levels in cell culture supernatants or cell-free lung lysates were obtained using L929 ISRE-luciferase reporter cells as previously described (Woodward et al., 2010). Briefly, luciferase reporter cells were seeded in a 96-well plate 24 h prior to incubation with supernatants or lung lysates for 6-8 h. Luciferase reporter activity was measured using the Luciferase Reporter Assay (Promega) according to the manufacturer’s instructions. Cytokines from cell culture supernatant or lung lysates were measured using DuoSet or Quantikine ELISA Kits (R&D Systems) following manufacturer’s protocols.

### SILAC labelling and di-glycine peptide sample preparation

RAW 264.7 mouse macrophages were labelled for 15 days in SILAC DMEM (Thermo Scientific #89985) supplemented with heavy or light labeled arginine (Thermo Scientific #89989 and #89990), heavy or light labeled lysine (Thermo Scientific #89987 and #89988), 10% dialyzed FBS (Thermo Scientific #88212), and 20 mM HEPES buffer. Mock and bacterial infections (m.o.i of 10) were carried out in the described SILAC media supplemented with 10% dialyzed horse serum. At the indicated time points, cells were fixed for 10 minutes in 100% methanol and then washed three times with PBS. Cell were lysed in 8M urea, 150 mM NaCl, 100 mM Tris pH 8.0 and protease inhibitor cocktail tablets (Roche #04693124001), sonicated two times at seven watts (Sonics and Materials Inc. model VC 130), and protein concentrations were measured using a micro BCA kit (Thermo #23235). Five mg of light-labeled lysate was mixed with five mg of heavy-labeled lysate, and the mixture was treated with 5 mM TCEP for 20 minutes at 60°C, chilled on ice for 10 minutes, then treated with 10 mM iodoacetamide at 25°C for 15 minutes in the dark. Lysates were then diluted to 2M urea with 100 mM Tris pH 8.0 prior to trypsin digestion, and 100 μg trypsin (Promega v5280) was added and incubated for 16 hours at 25°C. Peptides were purified over a c18 column (Waters Wat023590) according to manufacturer’s instructions and lyophilized for 48 hours.

### Immunoprecipitation of di-glycine peptides

Lyophilized peptides were resuspended in 1 mL IAP buffer (50 mM MOPS (pH 7.4), 10 mM Na_2_HPO_4_, 50 mM NaCl) and sonicated for 30 minutes at 4°C (Diagenode Bioruptor). Anti-di-glycine peptide beads (Cell Signalling #5562) were washed twice with IAP buffer, and incubated with resuspended peptides for 90 minutes at 4°C. Beads were washed three times with IAP buffer, twice with water, and eluted with 0.15% trifluoroacetic acid. The eluted peptides were then desalted using the Stage-tip method and the samples were dried.

### Preparation of samples for global ubiquitylation and phosphorylation analysis

Cells were lysed in a buffer containing 8M urea, 50 mM ammonium bicarbonate, 150 mM NaCl, and PhosStop and Complete-EDTA free phosphatase and protease inhibitors (Roche) and sonicated to sheer membranes and DNA. Protein concentrations were measured by a Bradford assay. For SILAC analyses, light and heavy cells for each comparison were combined at equal protein concentrations. Lysates were reduced by the addition of 4 mM TCEP (Sigma) for 30 minutes at room temperature, disulfide bonds were alkylated with 10 mM iodoacetamide (Sigma) for 30 minutes in the dark at room temperature, and excess iodoacetamide was quenched with 20 mM DTT (Sigma). Lysates were diluted 1:4 in 50 mM ammonium bicarbonate and trypsin was added at a 1:100 enzyme:substrate ratio. Lysates were digested for 18 hours at room temperature with rotation. Digested lysates were acidified with 0.1% TFA and peptides concentrated on Sep-Pak C18 solid phase extraction columns (Waters). For ubiquitin remnant analysis, an amount of digested lysate equivalent to 10 mg of protein was subjected to ubiquitin remnant immunoprecipitation according to the manufacturer’s protocol (Cell Signaling Technologies). For phosphorylation analysis, an amount of digested lysate equivalent to 1 mg of protein was lyophilized and then resuspended in a buffer containing 75% ACN with 0.1% TFA. Peptides were incubated with Fe^3+^-immobilized metal affinity chromatography (IMAC) beads, washed with the same resuspension buffer, and then phosphopeptides were eluted with 500 mM HK_2_PO_4_. For both ubiquitin remnant-enriched and phosphopeptide-enriched samples, the purified material was desalted using homemade C18 STAGE tips, evaporated to dryness, and then resuspended in 0.1% formic acid for mass spectrometry analysis (Rappsilber et al., 2007).

### Mass spectrometry analysis

All samples were analyzed on a Thermo Scientific LTQ Orbitrap Elite mass spectrometry system equipped with an Easy nLC 1000 uHPLC system interfaced with the mass spectrometry via a Nanoflex II nanoelectrospray source. Samples were injected onto a C18 reverse phase capillary column (75 μm inner diameter × 25 cm, packed with 1.9 μm Reprosil Pur C18-AQ particles). Peptides were then separated by an organic gradient from 5% to 30% in 0.1% formic acid at a flow rate of 300 nL/min. Ubiquitylation samples were separated over a 112 min gradient, and phosphorylation samples were separated over a 172 min gradient. The mass spectrometry collected data in a data-dependent fashion, collecting one full scan in the Orbitrap at 120,000 resolution followed by 20 collision-induced dissociation MS/MS scans in the dual linear ion trap for the 20 most intense peaks from the full scan. Charge state screening was enabled to reject MS/MS analysis of singly charged species or species for which a charge could not be assigned. Dynamic exclusion was enabled with a repeat count of 1, a duration of 30 s, and with mass widths of +/− 10 ppm surrounding all isotope peaks in an isotopic envelope. Raw mass spectrometry data were matched to peptide/protein sequences and SILAC ratios extracted using the MaxQuant platform (version 1.6.8) (Pereira et al., 2019). Data were searched against the SwissProt mouse UniProt sequence database (downloaded on 10/13/16). Peptides were required to have full trypsin specificity and up to two missed cleavages were allowed per peptide. Variable modifications were allowed for oxidation of methionine residues, acetylation of protein N-termini, diglycine-modified lysine residues for ubiquitylation analysis, and phosphorylation of serine, threonine, or tyrosine resides for phosphorylation analysis. The match between runs feature was enabled for +/− 2 min. All other MaxQuant settings were left at the default parameters. Protein significance analysis was performed using MSstats (Choi et al., 2014).

### Gene ontology enrichment analysis

The Metascape tool was used to identify significantly enriched gene ontology (GO) terms for molecular function, biological processes, and cellular compartments. Non-unique site identifications were removed prior to enrichment analysis. The gene list was comprised of UniProt accessions for differentially abundant ubiquitylation or phosphorylation sites using the criteria |log_2_fold-change| > 1.0 and adjusted p-value < 0.05. The background gene list was comprised of UniProt accession for all ubiquitylation or phosphorylation sites detected and quantified regardless of log_2_fold-change or adjusted p-values.

### PhosFate analysis

For PhosFate analysis, mouse phosphorylation sites positions were first converted to human phosphorylation site position. For each mouse phosphorylation site, the corresponding ortholog in human was mapped using the BioMart ortholog table (Smedley et al., 2009). The Blast algorithm was then used to align the mouse and human protein sequences. Alignments were processed in Perl and the mouse site was converted to the aligned human site if it was the same residue in both mouse and human. Human-converted phosphorylation log_2_fold-change profiles were then uploaded to PhosFate for kinase activity and condition correlation analysis.

### Statistics

Statistical analysis of data was performed using GrahPad Prism. Two-tailed unpaired Student’s t-tests were used for the analysis of bacterial and viral growth assays along with mouse CFU assays and cytokine measurements. Statistical analysis of the combined set of biological replicates was performed using an in-house computational pipeline, built around the MSstats bioconductor package. Briefly, after median-centering the log2-scaled intensity distributions across all replicates, a mixed or fixed effect model was fitted for every protein or modification site to account for systematic sources of variation when computing the significance test. Adjustments were made for all test results for multiple testing by applying a Benjamini-Hochberg correction on the resulting p-values and applied a composite threshold (p<0.05 and log2-fold-change>1 or <−1) to determine the set of significantly changing proteins or sites.

## SUPPLEMENTAL INFORMATION

**Table S1. Global ubiquitylation changes in response to *Lm*, *St*, and *Mtb* infection, Related to Figure 1**.

(A) MSstats results for global ubiquitylation changes.

(B) Extracted ion intensities for modified peptides detected.

(C) Description of column headers for tabs (A) and (B)

**Table S2. Global phosphorylation changes in response to *Lm*, *St*, and *Mtb* infection, Related to Figure 2**.

(A) MSstats results for global phosphorylation changes.

(B) MSstats results for global phosphorylation changes with mouse site accessions converted to human site accessions.

(C) Extracted ion intensities for modified peptides detected.

(D) Description of column headers for tabs (A) and (B)

**Table S3. Global ubiquitylation changes in response to wild-type and Δesx1 *Mtb* infection**.

(A) MSstats results for global ubiquitylation changes.

(B) Extracted ion intensities for modified peptides detected.

(C) Description of column headers for tabs (A) and (B)

**Supplementary Figure S1.**
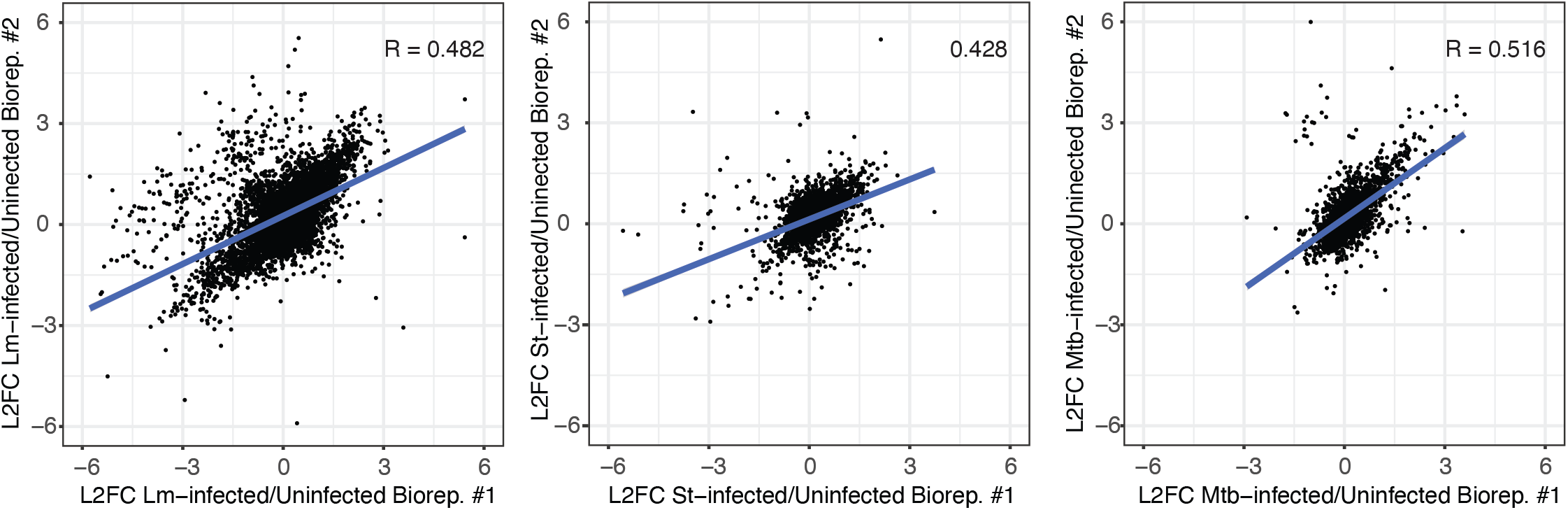
Log_2_fold-change scatter plots comparing biological replicate global ubiquitination analyses.

**Supplementary Figure S2.**
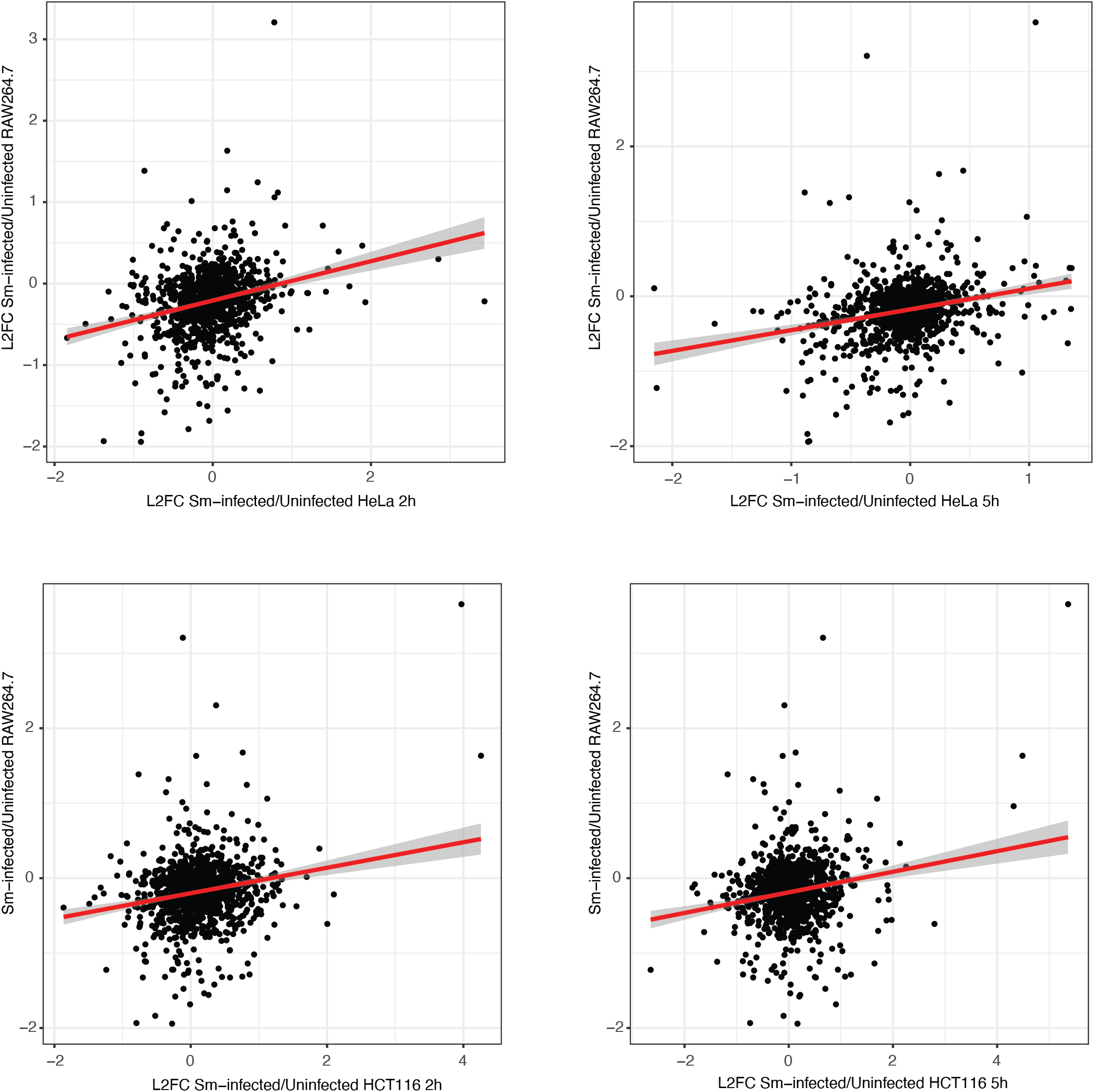
Log_2_fold-change scatter plots comparing *Salmonella enterica* serovar Typhimurium-infected RAW264.7 cells, HeLa cells, and HCT116 cells.

**Supplementary Figure S3.**
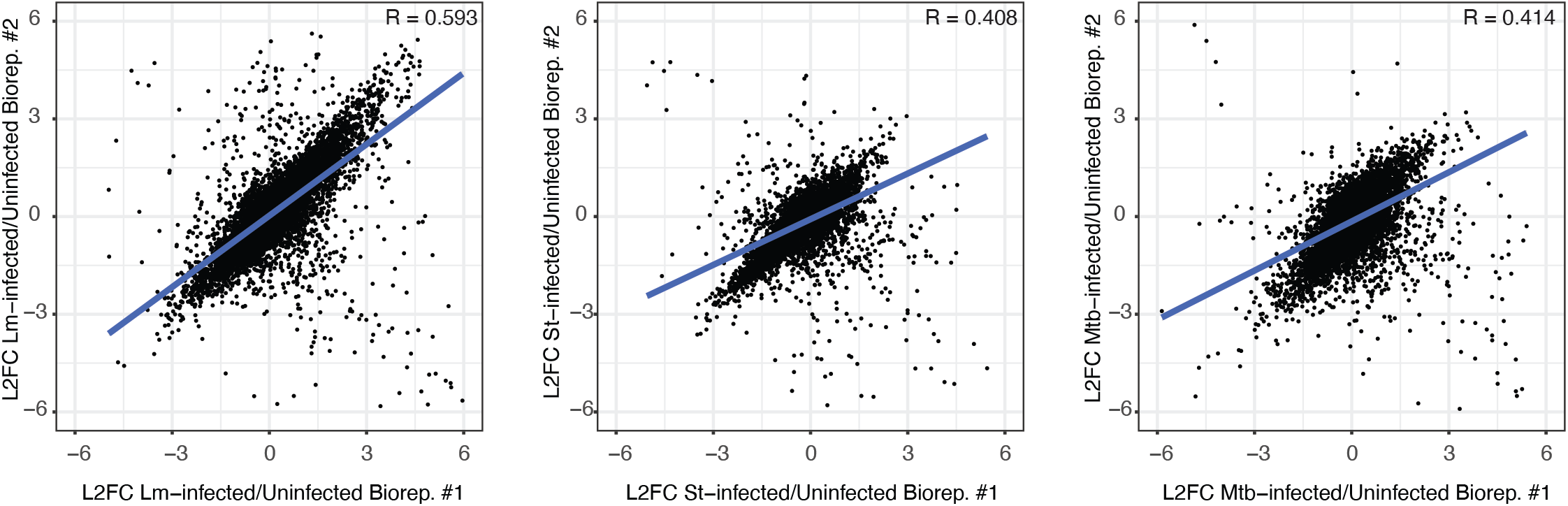
Log_2_fold-change scatter plots comparing biological replicate global phosphorylation analyses.

**Supplementary Figure S4.**
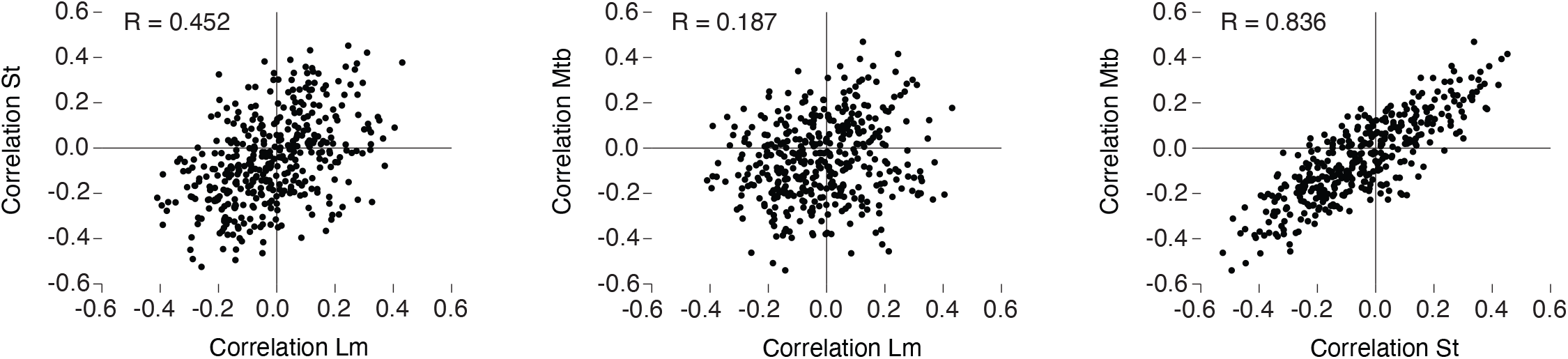
Scatter plots comparing correlation coefficients to existing phosphoproteomics data in the PhosFate database for different IBP infections.

**Supplementary Figure S5.**
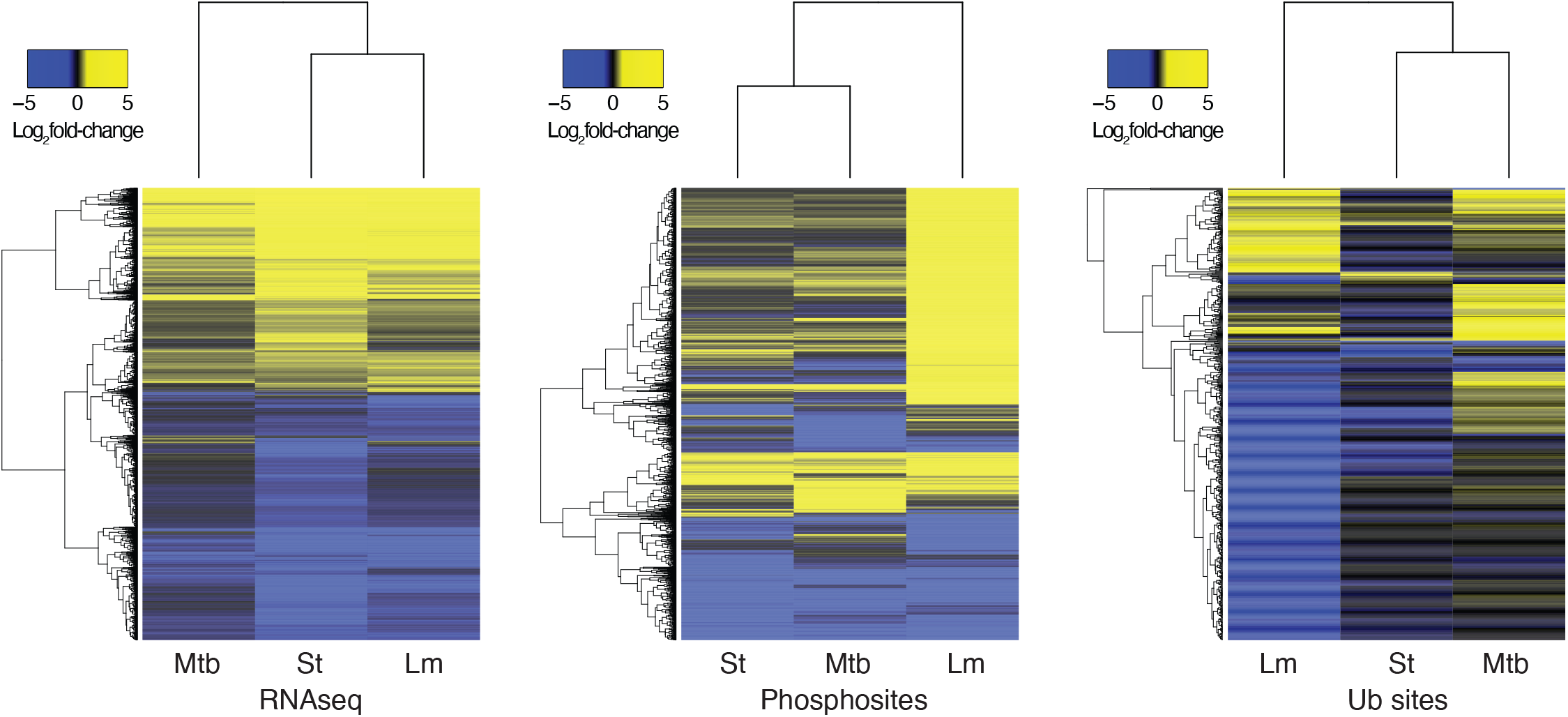
Hierarchically clustered heatmaps of log_2_fold-changes for global transcriptomics, phosphorylation, and ubiquitination analyses.

**Supplementary Figure S6.**
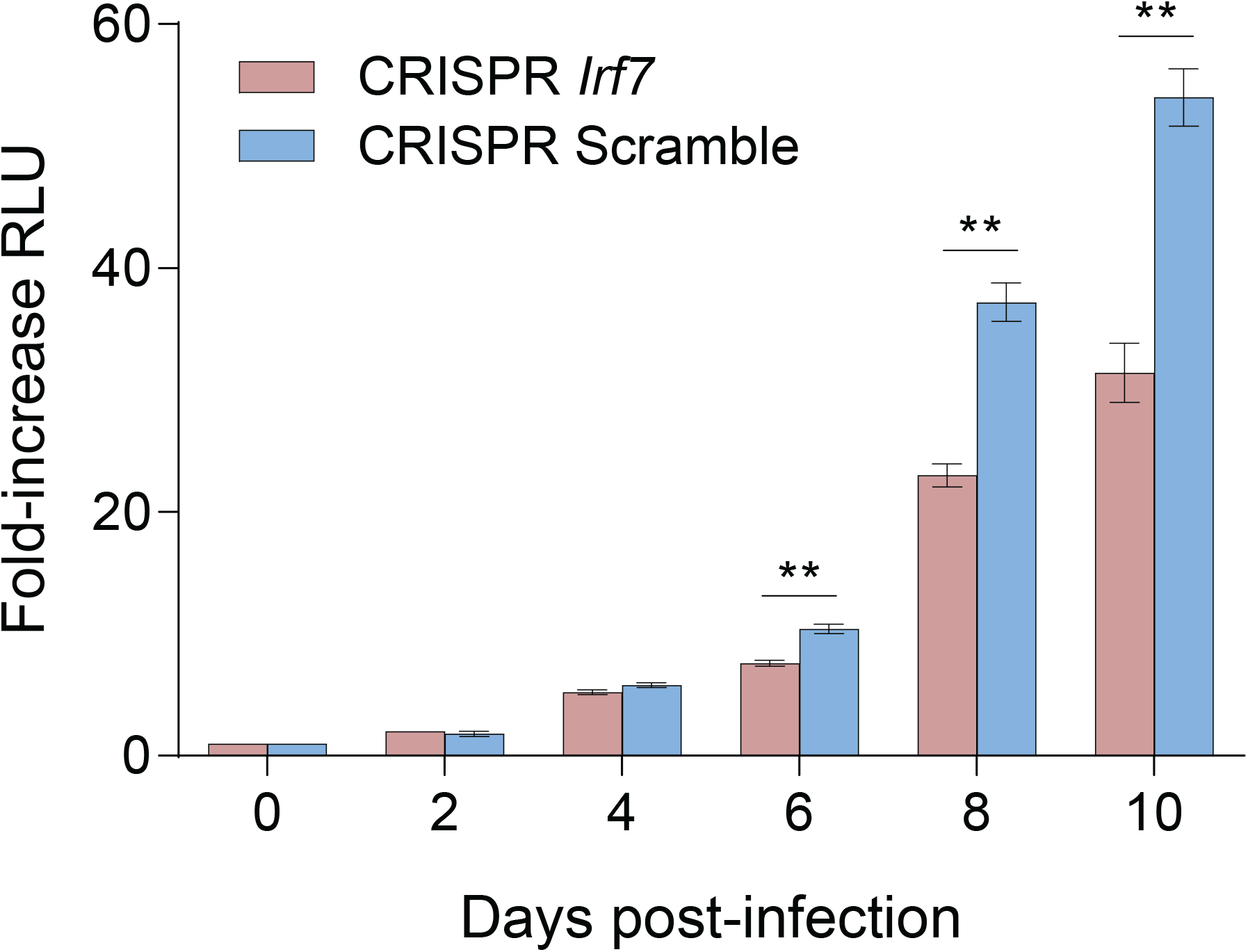
Related to Figure 6. *M. tuberculosis* growth in CRISPR mutagenized RAW 264.7 cells. Bacterial growth assay in CRISPR/Cas9 Irf7 mutant RAW 264.7 cells. CRISPR/Cas9-induced mutagenesis in the Irf7 locus or non-targeting (scramble) DNA. Lentiviral transduced RAW 264.7 cells were selected using antibiotics, followed by isolation of single cell clones.

**Supplementary Figure S7.**
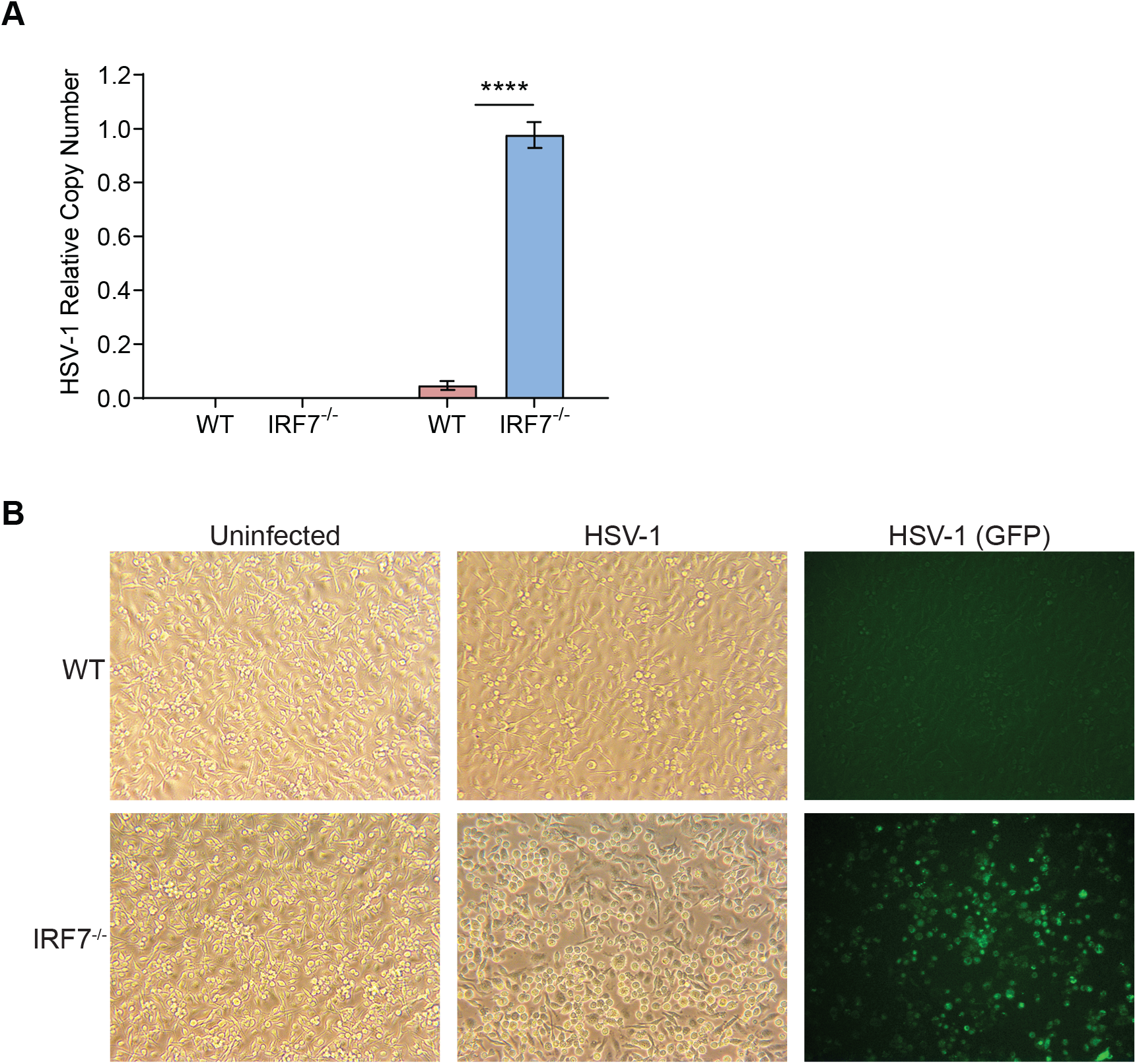
Related to Figure 5. *Irf7^−/−^* bone marrow-derived macrophage sensitivity to HSV-1 infection. (A) BMMs were infected with HSV-1 for 24 h and viral copy number from supernatant were enumerated by RT-qPCR. (B) Representative brightfield and fluorescent images of BMMs infected with HSV-1-eGFP.

**Supplementary Figure S8.**
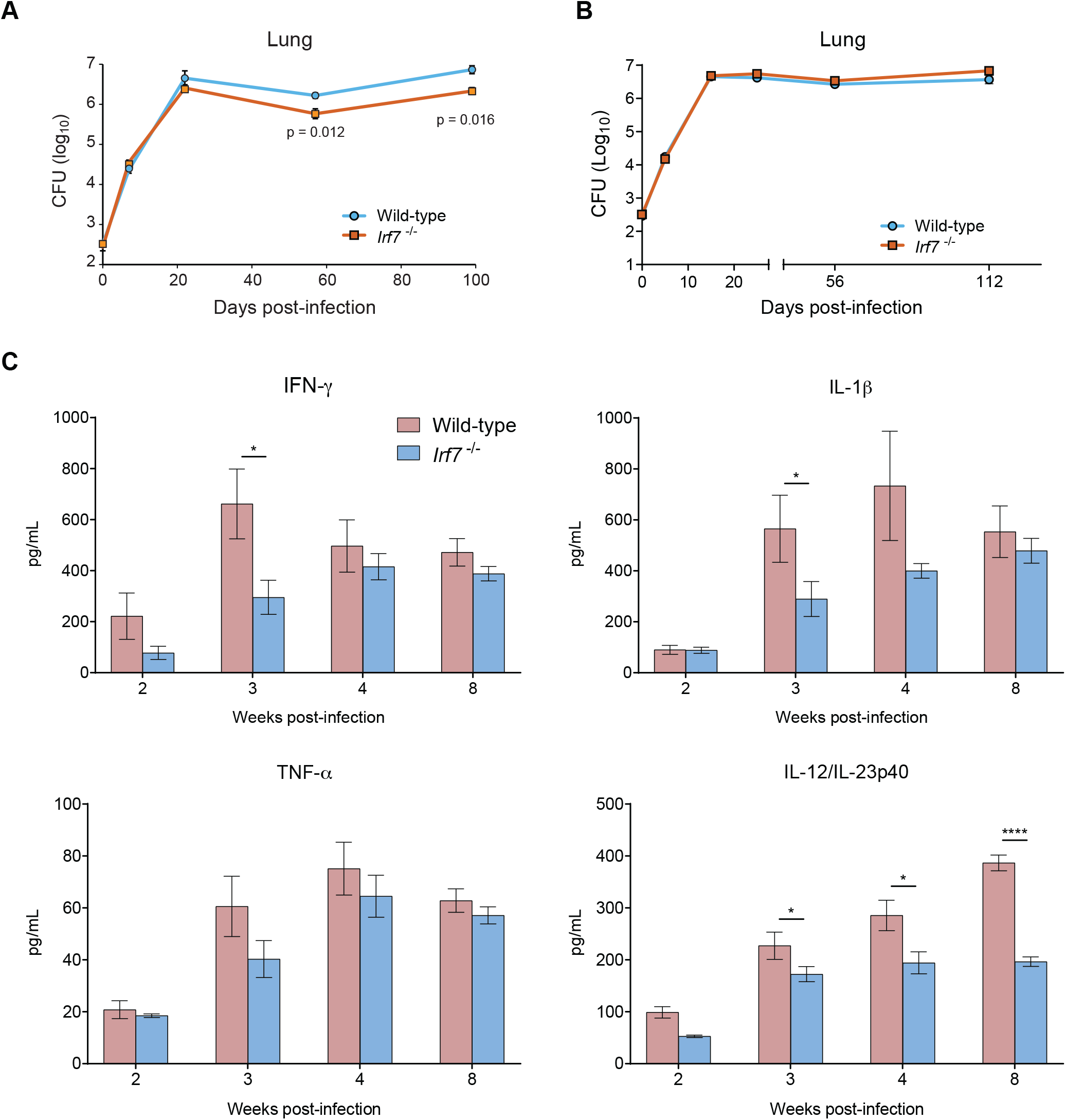
Related to Figure 6. *in vivo M. tuberculosis* infections performed at two different facilities, UCSF and UCB. (A) Wild-type and *Irf7*^−/−^ B6 mice (male and female) were infected with *M. tuberculosis* via aerosol route using Madison chamber device at UCSF. (B) Wild-type and *Irf7*^−/−^ B6 mice (male and female) were infected with *M. tuberculosis* via aerosol route using the Glas-Col inhalation exposure system at UCB. At indicated time points, lungs were homogenized and plated on 7H10 plates, and CFU were enumerated 21 days after plating. Data shown is representative of three separate experiments. (C) Pulmonary inflammatory cytokines analyzed by ELISA from one of the three mouse infection experiments at UCB of wild-type and *Irf7^−/−^* mice infected with *M. tuberculosis*.

**Supplementary Figure S9.**
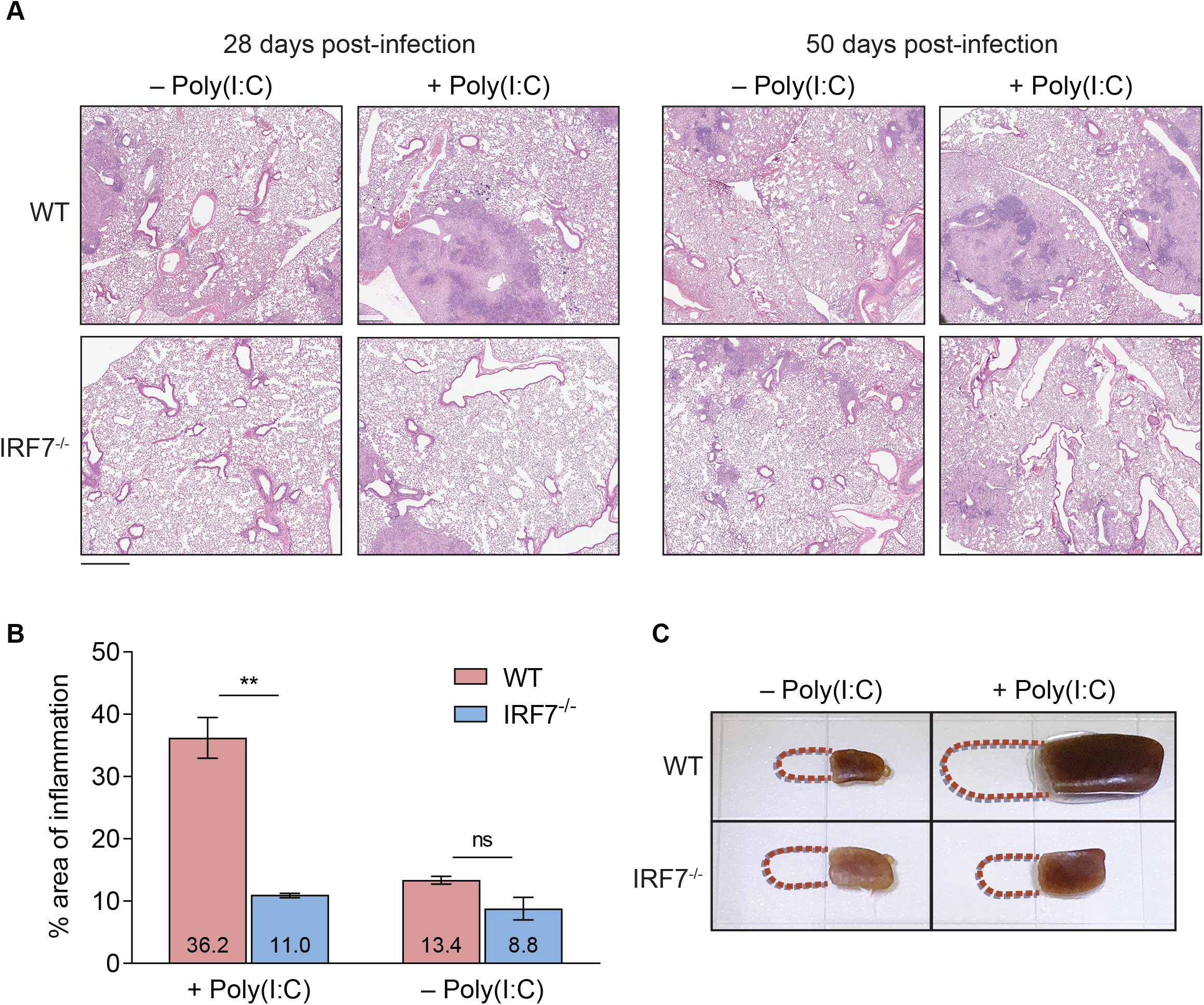
Related to Figure 6. Pathogenesis of *M. tuberculosis*-infected mice treated with poly(I:C). (A) Representative images of *M. tuberculosis*-infected mice lung tissues stained with H&E at indicated time points. (Scale baR=1 mm) (B) At 12 weeks post-infection, H&E stained lung tissues from *M. tuberculosis*-infected mice, either treated with poly(I:C) or PBS, were analyzed for lung inflammation. Data is ratio of total cumulative regions of inflamed lung sections/total area of lungs lobes analyzed. (C) Gross pathology of spleens from *M. tuberculosis*-infected mice treated with poly(I:C) or PBS at 12 weeks post-infection.

## REFERENCES

Antonelli, L.R.V., Gigliotti Rothfuchs, A., Gonçalves, R., Roffê, E., Cheever, A.W., Bafica, A., Salazar, A.M., Feng, C.G., and Sher, A. (2010). Intranasal Poly-IC treatment exacerbates tuberculosis in mice through the pulmonary recruitment of a pathogen-permissive monocyte/macrophage population. The Journal of Clinical Investigation 120, 1674–1682.

Beuzón, C.R., Méresse, S., Unsworth, K.E., Ruíz-Albert, J., Garvis, S., Waterman, S.R., Ryder, T.A., Boucrot, E., and Holden, D.W. (2000). Salmonella maintains the integrity of its intracellular vacuole through the action of SifA. Embo J. 19, 3235–3249.

Birmingham, C.L., Smith, A.C., Bakowski, M.A., Yoshimori, T., and Brumell, J.H. (2006). Autophagy controls Salmonella infection in response to damage to the Salmonella-containing vacuole. J Biol Chem 281, 11374–11383.

Braverman, J., Sogi, K.M., Benjamin, D., Nomura, D.K., and Stanley, S.A. (2016). HIF-1α Is an Essential Mediator of IFN-γ-Dependent Immunity to Mycobacterium tuberculosis. J Immunol 197, 1287–1297.

Cheng, S., Sun, C., Lao, W., and Kang, H. (2019). Association of VAMP8 rs1010 Polymorphism with Host Susceptibility to Pulmonary Tuberculosis in a Chinese Han Population. Genet Test Mol Biomarkers 23, 299–303.

Choi, M., Chang, C.-Y., Clough, T., Broudy, D., Killeen, T., MacLean, B., and Vitek, O. (2014). MSstats: an R package for statistical analysis of quantitative mass spectrometry-based proteomic experiments. Bioinformatics 30, 2524–2526.

Choy, A., Dancourt, J., Mugo, B., O’Connor, T.J., Isberg, R.R., Melia, T.J., and Roy, C.R. (2012). The Legionella effector RavZ inhibits host autophagy through irreversible Atg8 deconjugation. Science 338, 1072–1076.

Cumming, B.M., Rahman, M.A., Lamprecht, D.A., Rohde, K.H., Saini, V., Adamson, J.H., Russell, D.G., and Steyn, A.J.C. (2017). Mycobacterium tuberculosis arrests host cycle at the G1/S transition to establish long term infection. PLoS Pathog 13, e1006389.

De Luca, M., Cogli, L., Progida, C., Nisi, V., Pascolutti, R., Sigismund, S., Di Fiore, P.P., and Bucci, C. (2014). RILP regulates vacuolar ATPase through interaction with the V1G1 subunit. J Cell Sci 127, 2697–2708.

Deng, L., Wang, C., Spencer, E., Yang, L., Braun, A., You, J., Slaughter, C., Pickart, C., and Chen, Z.J. (2000). Activation of the IkappaB kinase complex by TRAF6 requires a dimeric ubiquitin-conjugating enzyme complex and a unique polyubiquitin chain. Cell 103, 351–361.

Deretic, V., and Levine, B. (2018). Autophagy balances inflammation in innate immunity. Autophagy 14, 243–251.

Fei, L., and Xu, H. (2018). Role of MCM2-7 protein phosphorylation in human cancer cells. Cell Biosci 8, 43.

Fiskin, E., Bionda, T., Dikic, I., and Behrends, C. (2016). Global Analysis of Host and Bacterial Ubiquitinome in Response to Salmonella Typhimurium Infection. Mol Cell 62, 967–981.

Galván-Peña, S., and O’Neill, L.A.J. (2014). Metabolic reprograming in macrophage polarization. Front. Immunol. 5, 420.

Gerarduzzi, C., He, Q., Antoniou, J., and Di Battista, J.A. (2014). Quantitative phosphoproteomic analysis of signaling downstream of the prostaglandin e2/g-protein coupled receptor in human synovial fibroblasts: potential antifibrotic networks. J Proteome Res 13, 5262–5280.

Gröschel, M.I., Sayes, F., Siméone, R., Majlessi, L., and Brosch, R. (2016). ESX secretion systems: mycobacterial evolution to counter host immunity. Nat Rev Microbiol 14, 677–691.

Härtlova, A., Herbst, S., Peltier, J., Rodgers, A., Bilkei-Gorzo, O., Fearns, A., Dill, B.D., Lee, H., Flynn, R., Cowley, S.A., et al. (2018). LRRK2 is a negative regulator of Mycobacterium tuberculosis phagosome maturation in macrophages. Embo J 37, e98694.

Herhaus, L., and Dikic, I. (2015). Expanding the ubiquitin code through post-translational modification. EMBO Reports 16, 1071–1083.

Honda, K., and Littman, D.R. (2016). The microbiota in adaptive immune homeostasis and disease. Nature 535, 75–84.

Honda, K., Yanai, H., Negishi, H., Asagiri, M., Sato, M., Mizutani, T., Shimada, N., Ohba, Y., Takaoka, A., Yoshida, N., et al. (2005). IRF-7 is the master regulator of type-I interferon-dependent immune responses. Nature 434, 772–777.

Huang, J., and Brumell, J.H. (2014). Bacteria-autophagy interplay: a battle for survival. Nat Rev Microbiol 12, 101–114.

Iwata, H., Goettsch, C., Sharma, A., Ricchiuto, P., Goh, W.W.B., Halu, A., Yamada, I., Yoshida, H., Hara, T., Wei, M., et al. (2016). PARP9 and PARP14 cross-regulate macrophage activation via STAT1 ADP-ribosylation. Nat Comms 7, 12849.

Ji, D.X., Yamashiro, L.H., Chen, K.J., Mukaida, N., Kramnik, I., Darwin, K.H., and Vance, R.E. (2019). Type I interferon-driven susceptibility to Mycobacterium tuberculosis is mediated by IL-1Ra. Nat Microbiol 4, 2128–2135.

Kamaruzzaman, N.F., Kendall, S., and Good, L. (2017). Targeting the hard to reach: challenges and novel strategies in the treatment of intracellular bacterial infections. Br. J. Pharmacol. 174, 2225–2236.

Karg, E., Smets, M., Ryan, J., Forné, I., Qin, W., Mulholland, C.B., Kalideris, G., Imhof, A., Bultmann, S., and Leonhardt, H. (2017). Ubiquitome Analysis Reveals PCNA-Associated Factor 15 (PAF15) as a Specific Ubiquitination Target of UHRF1 in Embryonic Stem Cells. J Mol Biol 429, 3814–3824.

Kim, W., Bennett, E.J., Huttlin, E.L., Guo, A., Li, J., Possemato, A., Sowa, M.E., Rad, R., Rush, J., Comb, M.J., et al. (2011). Systematic and quantitative assessment of the ubiquitin-modified proteome. Mol Cell 44, 325–340.

Kotewicz, K.M., Ramabhadran, V., Sjoblom, N., Vogel, J.P., Haenssler, E., Zhang, M., Behringer, J., Scheck, R.A., and Isberg, R.R. (2017). A Single Legionella Effector Catalyzes a Multistep Ubiquitination Pathway to Rearrange Tubular Endoplasmic Reticulum for Replication. Cell Host & Microbe 21, 169–181.

Lamothe, B., Besse, A., Campos, A.D., Webster, W.K., Wu, H., and Darnay, B.G. (2007). Site-specific Lys-63-linked tumor necrosis factor receptor-associated factor 6 auto-ubiquitination is a critical determinant of I kappa B kinase activation. J. Biol. Chem. 282, 4102–4112.

Laurell, E., Beck, K., Krupina, K., Theerthagiri, G., Bodenmiller, B., Horvath, P., Aebersold, R., Antonin, W., and Kutay, U. (2011). Phosphorylation of Nup98 by multiple kinases is crucial for NPC disassembly during mitotic entry. Cell 144, 539–550.

Lee, M.S., Kim, B., Oh, G.T., and Kim, Y.-J. (2013). OASL1 inhibits translation of the type I interferon-regulating transcription factor IRF7. Nat Immunol 14, 346–355.

Liang, S., Wang, F., Bao, C., Han, J., Guo, Y., Liu, F., and Zhang, Y. (2019). BAG2 ameliorates endoplasmic reticulum stress-induced cell apoptosis in Mycobacterium tuberculosis-infected macrophages through selective autophagy. Autophagy 195, 1–15.

Linder, M.I., Köhler, M., Boersema, P., Weberruss, M., Wandke, C., Marino, J., Ashiono, C., Picotti, P., Antonin, W., and Kutay, U. (2017). Mitotic Disassembly of Nuclear Pore Complexes Involves CDK1- and PLK1-Mediated Phosphorylation of Key Interconnecting Nucleoporins. Dev Cell 43, 141–156.e147.

Manca, C., Tsenova, L., Bergtold, A., Freeman, S., Tovey, M., Musser, J.M., Barry, C.E.3., Freedman, V.H., and Kaplan, G. (2001). Virulence of a Mycobacterium tuberculosis clinical isolate in mice is determined by failure to induce Th1 type immunity and is associated with induction of IFN-alpha /beta. Proc. Natl. Acad. Sci. U.S.a. 98, 5752–5757.

Manzanillo, P.S., Ayres, J.S., Watson, R.O., Collins, A.C., Souza, G., Rae, C.S., Schneider, D.S., Nakamura, K., Shiloh, M.U., and Cox, J.S. (2013). The ubiquitin ligase parkin mediates resistance to intracellular pathogens. Nature 501, 512–516.

Manzanillo, P.S., Shiloh, M.U., Portnoy, D.A., and Cox, J.S. (2012). Mycobacterium tuberculosis activates the DNA-dependent cytosolic surveillance pathway within macrophages. Cell Host & Microbe 11, 469–480.

Martinez, A., Lectez, B., Ramirez, J., Popp, O., Sutherland, J.D., Urbé, S., Dittmar, G., Clague, M.J., and Mayor, U. (2017). Quantitative proteomic analysis of Parkin substrates in Drosophila neurons. 12, 29.

Mayer-Barber, K.D., Andrade, B.B., Oland, S.D., Amaral, E.P., Barber, D.L., Gonzales, J., Derrick, S.C., Shi, R., Kumar, N.P., Wei, W., et al. (2014). Host-directed therapy of tuberculosis based on interleukin-1 and type I interferon crosstalk. Nature 511, 99–103.

McCormack, R.M., de Armas, L.R., Shiratsuchi, M., Fiorentino, D.G., Olsson, M.L., Lichtenheld, M.G., Morales, A., Lyapichev, K., Gonzalez, L.E., Strbo, N., et al. (2015a). Perforin-2 is essential for intracellular defense of parenchymal cells and phagocytes against pathogenic bacteria. Elife 4, 28039.

McCormack, R.M., Lyapichev, K., Olsson, M.L., Podack, E.R., and Munson, G.P. (2015b). Enteric pathogens deploy cell cycle inhibiting factors to block the bactericidal activity of Perforin-2. Elife 4, 263.

McCormack, R.M., Szymanski, E.P., Hsu, A.P., Perez, E., Olivier, K.N., Fisher, E., Goodhew, E.B., Podack, E.R., and Holland, S.M. (2017). MPEG1/perforin-2 mutations in human pulmonary nontuberculous mycobacterial infections. 2.

Medzhitov, R. (2008). Origin and physiological roles of inflammation. Nature 454, 428–435.

Medzhitov, R., and Horng, T. (2009). Transcriptional control of the inflammatory response. Nat. Rev. Immunol. 9, 692–703.

Mertins, P., Qiao, J.W., Patel, J., Udeshi, N.D., Clauser, K.R., Mani, D.R., Burgess, M.W., Gillette, M.A., Jaffe, J.D., and Carr, S.A. (2013). Integrated proteomic analysis of post-translational modifications by serial enrichment. Nat Meth 10, 634–637.

Mitchell, G., Chen, C., and Portnoy, D.A. (2016). Strategies Used by Bacteria to Grow in Macrophages. Microbiol Spectr 4.

Mitchell, G., Cheng, M.I., Chen, C., Nguyen, B.N., Whiteley, A.T., Kianian, S., Cox, J.S., Green, D.R., McDonald, K.L., and Portnoy, D.A. (2018). Listeria monocytogenes triggers noncanonical autophagy upon phagocytosis, but avoids subsequent growth-restricting xenophagy. Proc. Natl. Acad. Sci. U.S.a. 115, E210–E217.

Narayanan, L.A., and Edelmann, M.J. (2014). Ubiquitination as an efficient molecular strategy employed in salmonella infection. Front. Immunol. 5, 558.

Nau, G.J., Richmond, J.F.L., Schlesinger, A., Jennings, E.G., Lander, E.S., and Young, R.A. (2002). Human macrophage activation programs induced by bacterial pathogens. Proc Natl Acad Sci USA 99, 1503–1508.

Ochoa, D., Jonikas, M., Lawrence, R.T., Debs, El, B., Selkrig, J., Typas, A., Villén, J., Santos, S.D., and Beltrao, P. (2016). An atlas of human kinase regulation. Mol. Syst. Biol. 12, 888.

Ong, S.-E., Blagoev, B., Kratchmarova, I., Kristensen, D.B., Steen, H., Pandey, A., and Mann, M. (2002). Stable isotope labeling by amino acids in cell culture, SILAC, as a simple and accurate approach to expression proteomics. Mol. Cell Proteomics 1, 376–386.

Peng, H., Yang, J., Li, G., You, Q., Han, W., Li, T., Gao, D., Xie, X., Lee, B.-H., Du, J., et al. (2017). Ubiquitylation of p62/sequestosome1 activates its autophagy receptor function and controls selective autophagy upon ubiquitin stress. Cell Res 27, 657–674.

Penn, B.H., Netter, Z., Johnson, J.R., Dollen, Von, J., Jang, G.M., Johnson, T., Ohol, Y.M., Maher, C., Bell, S.L., Geiger, K., et al. (2018). An Mtb-Human Protein-Protein Interaction Map Identifies a Switch between Host Antiviral and Antibacterial Responses. Mol Cell 71, 637–648.e5.

Pereira, L.M.C., Bersano, P.R. de O., Moura, A. de A., and Lopes, M.D. (2019). First proteomic analysis of diestrus and anestrus canine oocytes at the germinal vesicle reveals candidate proteins involved in oocyte meiotic competence. Reprod. Domest. Anim. 54, 1532–1542.

Povlsen, L.K., Beli, P., Wagner, S.A., Poulsen, S.L., Sylvestersen, K.B., Poulsen, J.W., Nielsen, M.L., Bekker-Jensen, S., Mailand, N., and Choudhary, C. (2012). Systems-wide analysis of ubiquitylation dynamics reveals a key role for PAF15 ubiquitylation in DNA-damage bypass. Nature Cell Biology 14, 1089–1098.

Price, J.V., and Vance, R.E. (2014). The macrophage paradox. Immunity 41, 685–693.

Radoshevich, L., Impens, F., Ribet, D., Quereda, J.J., Nam Tham, T., Nahori, M.-A., Bierne, H., Dussurget, O., Pizarro-Cerdá, J., Knobeloch, K.-P., et al. (2015). ISG15 counteracts Listeria monocytogenes infection. Elife 4.

Rappsilber, J., Mann, M., and Ishihama, Y. (2007). Protocol for micro-purification, enrichment, pre-fractionation and storage of peptides for proteomics using StageTips. Nature Protocols 2, 1896–1906.

Sahan, A.Z., Hazra, T.K., and Das, S. (2018). The Pivotal Role of DNA Repair in Infection Mediated-Inflammation and Cancer. Front Microbiol 9, 663.

Schnupf, P., and Portnoy, D.A. (2007). Listeriolysin O: a phagosome-specific lysin. Microbes Infect 9, 1176–1187.

Siméone, R., Bobard, A., Lippmann, J., Bitter, W., Majlessi, L., Brosch, R., and Enninga, J. (2012). Phagosomal rupture by Mycobacterium tuberculosis results in toxicity and host cell death. PLoS Pathog 8, e1002507.

Smale, S.T., Tarakhovsky, A., and Natoli, G. (2014). Chromatin Contributions to the Regulation of Innate Immunity. Annu Rev Immunol 32, 489–511.

Smedley, D., Haider, S., Ballester, B., Holland, R., London, D., Thorisson, G., and Kasprzyk, A. (2009). BioMart--biological queries made easy. BMC Genomics 10, 22.

Sol, A., Lipo, E., de Jesús-Díaz, D.A., Murphy, C., Devereux, M., and Isberg, R.R. (2019). Legionella pneumophila translocated translation inhibitors are required for bacterial-induced host cell cycle arrest. Proceedings of the National Academy of Sciences 116, 3221–3228.

Stanley, S.A., Johndrow, J.E., Manzanillo, P., and Cox, J.S. (2007). The Type I IFN response to infection with Mycobacterium tuberculosis requires ESX-1-mediated secretion and contributes to pathogenesis. J Immunol 178, 3143–3152.

Stevens, J.M., Galyov, E.E., and Stevens, M.P. (2006). Actin-dependent movement of bacterial pathogens. Nat Rev Microbiol 4, 91–101.

Trevelin, S.C., Shah, A.M., Letters, G.L.I., 2020 (2020). Beyond Bacterial Killing: NADPH oxidase 2 is an immunomodulator. Elsevier.

Tripathi, S., Pohl, M.O., Zhou, Y., Rodriguez-Frandsen, A., Wang, G., Stein, D.A., Moulton, H.M., DeJesus, P., Che, J., Mulder, L.C.F., et al. (2015). Meta- and Orthogonal Integration of Influenza “OMICs” Data Defines a Role for UBR4 in Virus Budding. Cell Host & Microbe 18, 723–735.

van de Veerdonk, F.L., Netea, M.G., Dinarello, C.A., and Joosten, L.A.B. (2011). Inflammasome activation and IL-1β and IL-18 processing during infection. Trends Immunol. 32, 110–116.

Vance, R.E., Isberg, R.R., and Portnoy, D.A. (2009). Patterns of pathogenesis: discrimination of pathogenic and nonpathogenic microbes by the innate immune system. Cell Host & Microbe 6, 10–21.

Watson, R.O., Manzanillo, P.S., and Cox, J.S. (2012). Extracellular M. tuberculosis DNA targets bacteria for autophagy by activating the host DNA-sensing pathway. Cell 150, 803–815.

Woodward, J.J., Iavarone, A.T., and Portnoy, D.A. (2010). c-di-AMP secreted by intracellular Listeria monocytogenes activates a host type I interferon response. Science 328, 1703–1705.

Xia, Y., Liu, N., Xie, X., Bi, G., Ba, H., Li, L., Zhang, J., Deng, X., Yao, Y., Tang, Z., et al. (2019). The macrophage-specific V-ATPase subunit ATP6V0D2 restricts inflammasome activation and bacterial infection by facilitating autophagosome-lysosome fusion. Autophagy 15, 960–975.

Xu, G., Paige, J.S., and Jaffrey, S.R. (2010). Global analysis of lysine ubiquitination by ubiquitin remnant immunoaffinity profiling. Nat Biotechnol 28, 868–873.

Yang, C.-S., Jividen, K., Spencer, A., Dworak, N., Ni, L., Oostdyk, L.T., Chatterjee, M., Kuśmider, B., Reon, B., Parlak, M., et al. (2017). Ubiquitin Modification by the E3 Ligase/ADP-Ribosyltransferase Dtx3L/Parp9. Mol Cell 66, 503–516.e505.

Zhang, Y., Mao, D., Roswit, W.T., Jin, X., Patel, A.C., Patel, D.A., Agapov, E., Wang, Z., Tidwell, R.M., Atkinson, J.J., et al. (2015). PARP9-DTX3L ubiquitin ligase targets host histone H2BJ and viral 3C protease to enhance interferon signaling and control viral infection. Nat Immunol 16, 1215–1227.

Zhou, R., Yazdi, A.S., Menu, P., and Tschopp, J. (2011). A role for mitochondria in NLRP3 inflammasome activation. Nature 469, 221–225.

